# The nature of mitotic forces in epithelial monolayers

**DOI:** 10.1101/2020.11.16.378927

**Authors:** Vivek K. Gupta, Sungmin Nam, Jaclyn Camuglia, Judy Lisette Martin, Erin Nicole Sanders, Lucy Erin O’Brien, Adam C. Martin, Taeyoon Kim, Ovijit Chaudhuri

**Affiliations:** Department of Mechanical Engineering, Stanford University, CA, USA; Harvard John A. Paulson School of Engineering and Applied Sciences, Harvard University, MA, USA; Wyss Institute for Biologically Inspired Engineering, Cambridge, MA, USA; Department of Biology, Massachusetts Institute of Technology, MA, USA; Department of Molecular and Cellular Physiology, Stanford University, CA, USA; Weldon School of Biomedical Engineering, Purdue University, IN, USA

## Abstract

Epithelial cells undergo striking morphological changes during mitosis to ensure proper segregation of genetic and cytoplasmic materials. These morphological changes occur despite dividing cells being mechanically restricted by neighboring cells, indicating the need for extracellular force generation. While forces generated during mitotic rounding are well understood, forces generated after rounding remain unknown. Here, we identify two distinct stages of mitotic force generation that follow rounding: (1) protrusive forces along the mitotic axis that drive mitotic elongation, and (2) outward forces that facilitate post-mitotic re-spreading. Cytokinetic ring contraction of the mitotic cell, but not activity of neighboring cells, generates extracellular forces that propel mitotic elongation and also contribute to chromosome separation. Forces from mitotic elongation are observed in epithelia across many model organisms. Thus, forces from mitotic elongation represent a universal mechanism that powers mitosis in confining epithelia.

## Introduction

Epithelia are tightly packed sheets of cells that line the surfaces of organs and cavities. Cell division within epithelial tissues occurs continuously during development, homeostasis, and regeneration, and is critical for expanding tissues or replenishing cells lost due to extrusion, apoptosis, or injury ^1^. Dividing cells are surrounded by adjacent cells and the underlying basement membrane, creating a confining microenvironment (Fig. 1a). As cell division occurs typically within the plane of the monolayer, the extreme morphological changes that are required for cell division to progress normally must be accompanied by extracellular forces, or mitotic forces. Beyond driving cell division itself, mitotic forces have been implicated in contributing to distinct processes, including development, cell rearrangements, cell migration, epidermal stratification, and growth ^2–9^.

**Fig. 1.**
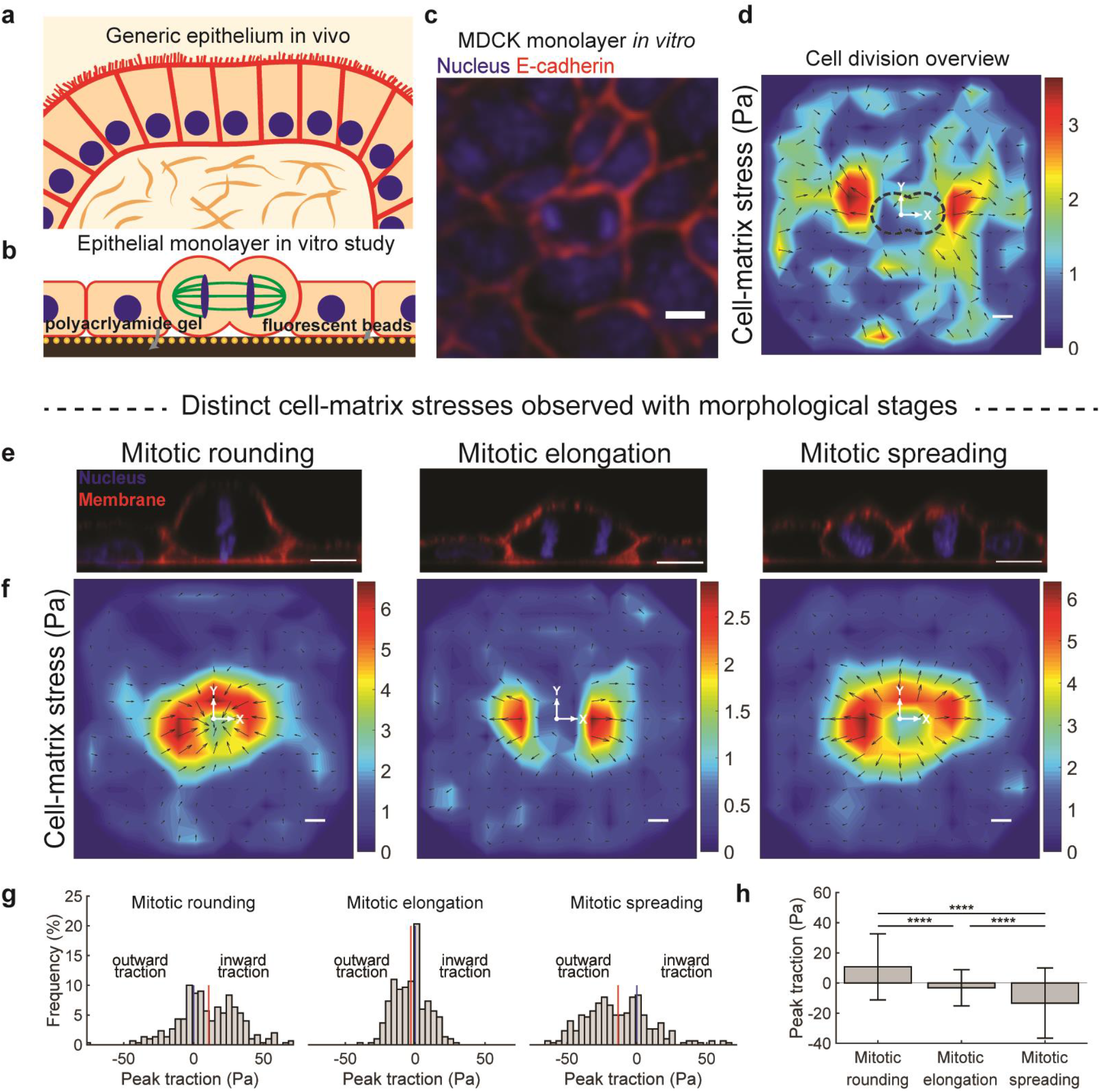
Cell division within epithelial monolayers is characterized by three distinct stages of cell-matrix and cell-cell stress: mitotic rounding, mitotic elongation, and post-mitotic re-spreading. **a**, Schematic of a generic epithelium *in vivo.* **b**, **c**, Experimental setup (**b**) and image (**c**) of MDCK epithelial monolayer grown on polyacrylamide gel substrate with embedded fluorescent beads. **d**, Net average change in cell-matrix stress over the entire cycle of cell division. **e**, Cross-sectional images of three morphological stages of cell division: mitotic rounding, mitotic elongation, and mitotic spreading. **f**-**h**, Average change in cell-matrix stress (**f**) and their distribution (**g**, **h**), during the three stages of division. Vertical red lines indicate means; blue line at 0 Pa; edge bins contain exceeding values as well. Multiple comparison test P value < 0.0001 (****). Scale bars, 10 μm.

While mitotic forces are critical for successful division completion, as well as various tissue-scale processes, the nature and origins of these forces remain unclear, beyond the well-studied process of mitotic rounding. In mitotic rounding, which occurs prior to metaphase, the dividing cell transitions to a rounded morphology by generating forces through a combination of actomyosin contractility and hydrostatic pressure ^10–14^. However, after mitotic rounding, the dividing cell continues to undergo morphological changes as it elongates along the mitotic axis and is cleaved at its center by a cytokinetic ring. Both of these processes are critical for proper segregation of chromosomes and other cytoplasmic materials and are required for successful completion of mitosis^15^. Further, the resulting rounded daughter cells then undergo spreading as they reintegrate into the monolayer to maintain proper barrier function. Though mitotic rounding in epithelia has been studied extensively, the extracellular forces associated during and after mitosis remain largely unexplored.

### Three stages of force generation during mitosis

Here, we investigated the forces accompanying cell division in epithelial monolayers. MDCK monolayers were grown on soft (~1kPa) polyacrylamide gels coated with type I collagen and embedded with fluorescent beads (Fig. 1b, c, and Supplementary Fig. 1). Cell division events were identified and traction force (TFM) and monolayer stress microscopy (MSM) were used to measure changes in cell-matrix and cell-cell stresses during cell division, respectively^16,17^. Because measured cell-matrix and cell-cell stresses are calculated based on forces applied by dynamic cell-matrix adhesions, many (>100) division events were averaged and aligned to isolate forces solely associated with cell division^18^. Through the entire course of cell-division, encompassing the time period over which a spread parent cell becomes two spread daughter cells, we found there to be a net outward cell-matrix stress (Fig. 1d, Supplementary Fig. 2a-i).

Based on morphological changes and measured stresses, cell division was categorized into three distinct chronological stages: mitotic rounding, mitotic elongation, and post-mitotic spreading (Fig. 1e). On average, net inward stresses develop during mitotic rounding of cells in relatively flat MDCK monolayers (Fig. 1e-h). However, mitotic rounding can be separated into two phases based on cell-matrix stresses, with net inward stresses generated during the initial phase, and net outward stresses occurring over the 6 minutes prior to metaphase (Supplementary Fig. 2j, k). Previously, more cuboidal or columnar epithelia have been associated only with net outward stresses during mitotic rounding, likely due to the different geometry of cells^10^. During mitotic elongation, encompassing elongation of the interpolar spindle at anaphase onset through cytokinesis completion, outward stresses develop, primarily along the mitotic axis. Finally, outward stresses develop in all directions as the daughter cells spread back onto the underlying substrate (Fig. 1f-h). Cell-cell stresses are consistent with cell-matrix stresses, with inward cell-matrix stresses associated with tensile cell-cell stresses, and outward cell-matrix stresses associated with compressive cell-cell stresses (Supplementary Fig. 2a-i). Measured stress values for mitotic elongation and mitotic spreading were comparable to those of mitotic rounding. We note that traction force and monolayer stress microscopy are likely to be underestimating the true value of mitotic stresses because cell division is occurring primarily at the center plane of the monolayer, but these techniques rely on measurements of substrate deformation that are made at the bottom monolayer plane. Indeed, previous measurements of mitotic rounding stresses with atomic force microscopy have reported stresses on the order of ~500 Pa^12^. Stresses during mitotic elongation were significantly more anisotropic and skewed along the division axis, with the average stress along the perpendicular axis less than 25% of the average stress along the mitotic axis, compared to mitotic rounding and spreading, in which perpendicular axis stresses were ~75% of mitotic axis stresses. Comparable results were found for MCF10A epithelial monolayers (Supplementary Fig. 3), and MDCK monolayers at varying densities (Supplementary Fig. 4). Single MDCK cells did not exhibit outward stresses during mitotic elongation (Supplementary Fig. 5). Thus, these analyses identified three distinct stages of extracellular force generation – mitotic rounding, mitotic elongation, and mitotic spreading – during cell division in epithelial monolayers.

### The dividing cell drives mitotic elongation

Next, we considered the origins of force generation in mitotic elongation and mitotic spreading, the two stages of extracellular mitotic force generation revealed in this study. While previous studies have not linked post-mitotic spreading of epithelial cells to outward force generation within monolayers, they do indicate that mitotic spreading is similar to general cell spreading, which is powered through the formation of cell-matrix adhesions and actin polymerization^19,20^. In contrast to the well-understood mechanisms underlying spreading, forces generated by mitotic elongation have only been studied in the context of single, isolated cancer cells embedded within inert alginate hydrogels, a context lacking neighboring cells and cell-matrix adhesions^15^.

Thus, we investigated the mechanisms underlying mitotic elongation in epithelial monolayers. We reasoned that there were three possible sources of the forces underlying mitotic elongation: (1) pulling forces from adjacent cells that cause elongation of the mitotic cell; (2) movement of adjacent cells away from the mitotic cells, which would open up space for mitotic elongation to occur as the cells de-adhere from the substrate; or, (3) the dividing cell itself pushes outward to generate space for mitotic elongation (Fig. 2a). Cell-cell adhesions within epithelial monolayers are expected to be under tension due to actomyosin contractility^21^, making it possible that tension transmitted across cell-cell adhesions along the mitotic axis pull the dividing cell apart. In this case similar patterns of neighboring cell arrangement relative to the mitotic cell would be expected, with the mitotic axis consistently aligned towards either adjacent cell edges or vertices. However, both these cases, and the case of a cell dividing against both a cell edge and vertex, were prevalent (Fig. 2b). Furthermore, the nuclei of adjacent cells were randomly distributed with respect to the dividing cell (Supplementary Fig. 6a, b), indicating that the mitotic axis orientation is independent of the arrangement of neighboring cells, making it unlikely that neighboring cells drive mitotic elongation. To further test this idea, the role of E-cadherin, a cell-cell adhesion protein that is known to transmit tension between cells to the actin-cytoskeleton, was examined by using cells over-expressing a mutant E-cadherin with a truncated extracellular domain that cannot transmit forces between cells^21,22^. In mosaic monolayers, division of truncated E-cadherin cells was accompanied by larger protrusive cell-matrix stresses, when compared to wild-type (WT) cells, indicating that the intracellular domain of E-cadherin was not necessary for the protrusive stresses (Fig. 2c, d, Supplementary Fig. 6c-e). Overall, these data establish that the forces of mitotic elongation do not arise from neighboring cells pulling on the dividing cell.

**Fig. 2.**
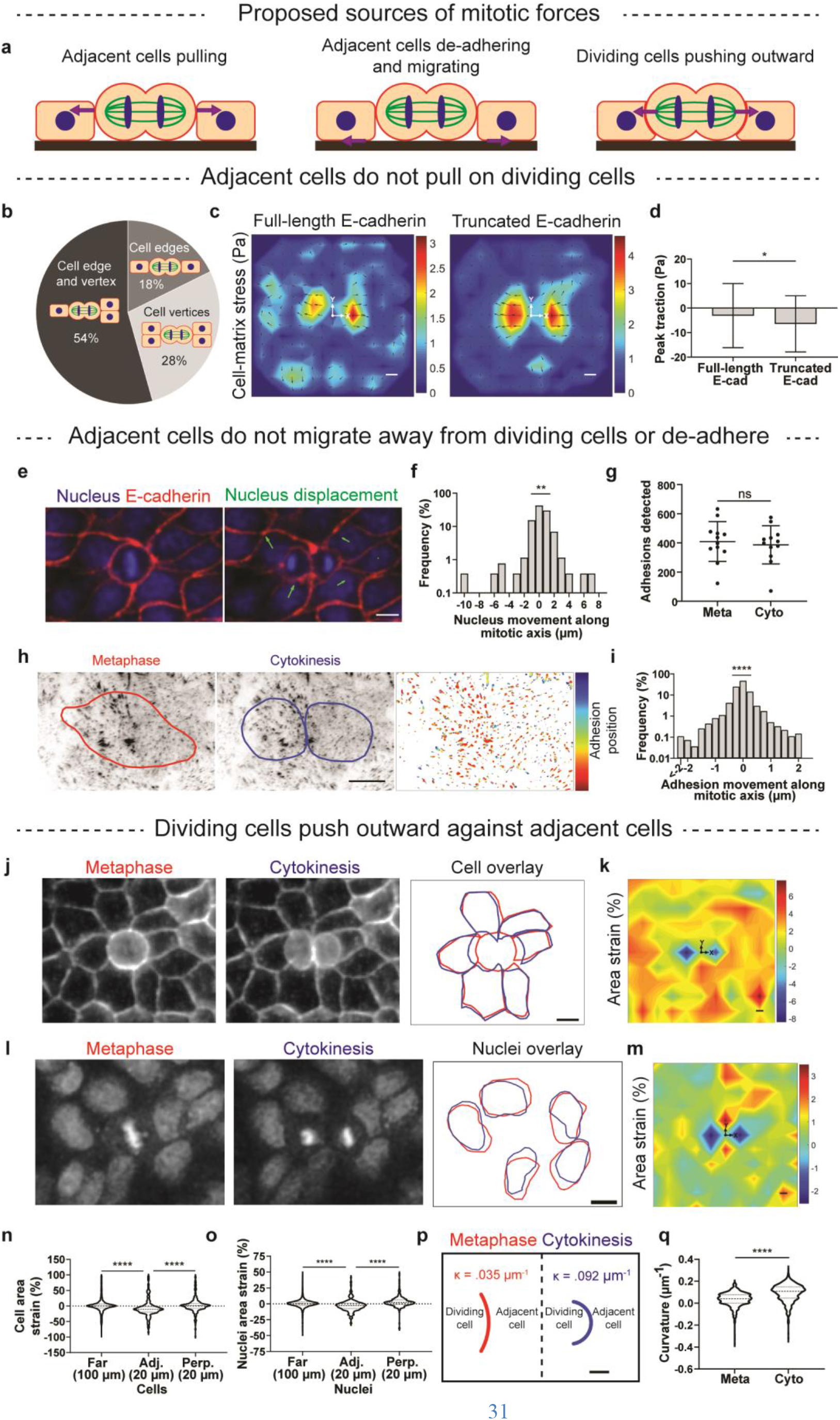
Forces generated during mitotic elongation originate from the dividing cell pushing outward against adjacent cells along the mitotic axis. **a**, Potential mechanisms that drive mitotic elongation include adjacent cells pulling, adjacent cells migrating, or the dividing cell pushing outward. **b**, Distribution of cells dividing against cell edges, cell vertices, or a cell edge and a vertex. **c**, **d**, Average change in cell-matrix stress during mitotic elongation generated with WT and truncated E-cadherin cells. Unpaired t-test. **e-f,** Movement of nuclei adjacent to a dividing cell. One-sample t-test. **g**, The number of detected adhesions at metaphase and cytokinesis. Paired t-test. **h**, **i**, Adhesions near the periphery of a dividing cell (**h**) and their movement (**i**). One-sample t-test. **j**-**m**, Images of adjacent cells (**j**) and nuclei (**k**) being deformed during mitotic elongation, with corresponding averaged area strain heat maps (**k**, **m**). **n**, **o**, Quantification of cell (**n**) and nuclei (**o**) area strain between adjacent, perpendicular, and neighboring cells. Tukey’s multiple comparison test. **p**, **q**, Schematic displaying average change in curvature (κ) of an adjacent cell (**p**) and quantification (**q**) of curvatures at metaphase and cytokinesis. Paired t-test. Scale bars, 10 μm. P values > 0.05 (n.s.), <0.05 (*), <0.01 (**), <0.001 (***), < 0.0001 (****).

Next, we considered the movement of cells adjacent to the dividing cell. Nuclei of adjacent cells were tracked as the dividing cell transitions from metaphase to cytokinesis. While motile, the neighboring nuclei were not consistently moving away from, or towards, the dividing cell (Fig. 2e, f). Cell adhesions were examined as release of neighboring cell adhesions near the mitotic cell could also create space for mitotic elongation to occur. MDCK cells stably expressing vinculin::GFP, a marker for focal adhesions, were imaged during division. Similar levels of adhesions were identified near the dividing cell at metaphase and cytokinesis (Fig. 2g). Furthermore, adhesions of adjacent cells did not consistently move away or toward the dividing cell (Fig. 2h, i). Taken together, these results support the conclusion that adjacent cells do not migrate away from dividing cells during mitosis or de-adhere.

With the role of the neighboring cells in driving mitotic elongation eliminated as a possibility, we directly investigated whether observed mitotic forces were originating from the dividing cell pushing outward against its neighbors. We imaged the cell membrane and nuclei of cells laying adjacent to the dividing cell along the division axis. As the dividing cell elongated, adjacent cells and their nuclei were deformed, exhibiting reduced cross-sectional areas at the center plane, compared to that of perpendicular or neighboring cells (Fig. 2j-o). Deformation of cells was only observed for cells along the division axis and synchronized with mitotic elongation. Further, the increased curvature (κ) of the neighboring cell edge along the mitotic axis indicates the application of protrusive forces by the dividing cell to the cell-edge (Fig. 2p, q). Taken together, these results indicate that mitotic elongation arises due to the dividing cell pushing outward.

### Force generation mechanisms for mitotic elongation

Next, we sought to determine how the dividing cell generates protrusive forces during mitotic elongation. A previous study found that cancer cells embedded within inert hydrogels exert protrusive forces through a combination of interpolar spindle elongation and cytokinetic ring contraction, which drives expansion along the mitotic axis due to conservation of volume (Fig. 3a)^15^. Consistent with the possibility of both these mechanisms contributing, cells elongate and generate compressive stresses through the entire course of mitotic elongation in epithelial monolayers (Fig. 3b, c). Therefore, we examined the contribution of these mechanisms to mitotic elongation.

**Fig. 3.**
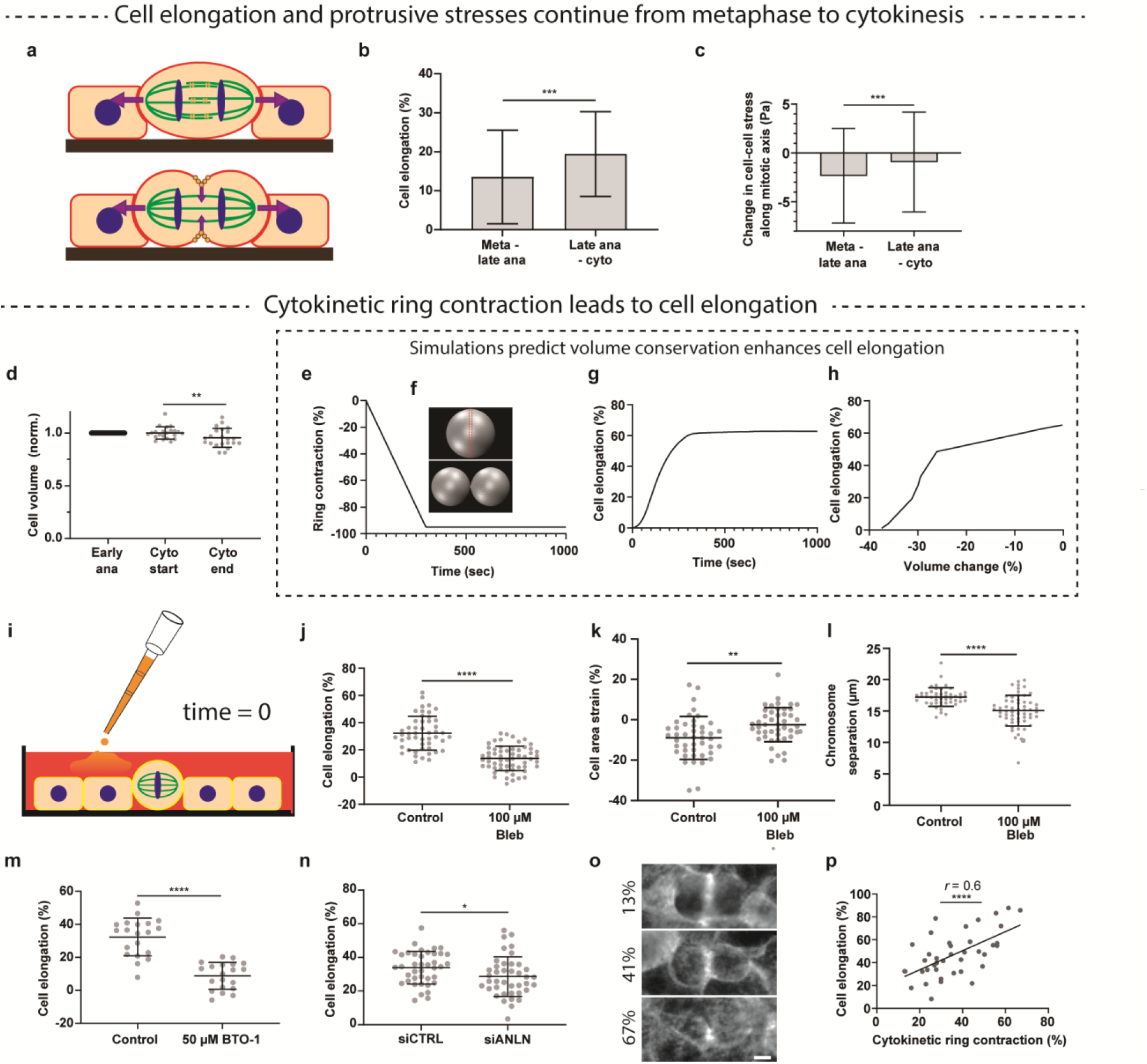
Forces for mitotic elongation are generated primarily from cytokinetic ring contraction, and to a lesser extent, interpolar spindle elongation. **a**, Forces for mitotic elongation can in principle be generated from interpolar spindle elongation and cytokinetic ring contraction. **b**, **c**, Cell elongation (**b**) and compressive stress generation (**c**) continue during the early (metaphase to late anaphase) and later (late anaphase to cytokinesis) stages of division. Unpaired t-test. **d**, Change in cell volume before and after cytokinesis. Paired t-test. **e**, **f**, Simulation of cytokinetic ring contraction progression (**e**), and corresponding computer model (**f**). **g**, Predicted cell elongation due to cytokinetic ring contraction for a ~5% cell volume reduction. **h**, Predicted cell elongation based on varying levels of cell volume reduction during division. (**I**) Schematic of inhibition experiments. **j**-**l**, Cell elongation (**j**), adjacent cell area strain (**k**), and chromosome separation (**l**) with blebbistatin treatment. **m**, **n**, Cell elongation with BTO-1 treatment (**m**) and anillin knockdown (**n**). Unpaired t-tests. **o**, Representative images of actin for anillin knockdown cells at maximum cytokinetic ring contraction. **p**, Correlation between max contractile ring contraction and cell elongation for anillin knockdown cells. Pearson’s *r*. Scale bars, 10 μm. P values > 0.05 (n.s.), < 0.05 (*), <0.01 (**), <0.001 (***), < 0.0001 (****).

We first determined the contribution of interpolar spindle elongation to protrusive force generation during mitotic elongation. During the early parts of elongation, kinesin motor proteins, including kinesin-5, push cross-linked interpolar microtubules apart, and these forces can potentially be transmitted to the neighboring cells via astral microtubules (Fig. 3a)^15^. However, interpolar spindle elongation was not significantly correlated with compressive stresses developed (Supplementary Fig. 7a, b). Further, inhibition of kinesin-5 with BRD9876^23^, applied to cells at prometaphase and onward (Fig. 3i), while reducing interpolar spindle elongation, only slightly diminished cell elongation (24% reduction) (Supplementary Fig. 7c, d). Next, laser ablation was used to sever the interpolar spindle of dividing cells during mitotic elongation. If the interpolar spindle elongation drove mitotic elongation, ablation of the structure should result in almost immediate retraction of the dividing cell’s chromatids and membrane. However, ablation resulted in chromatid and cell retraction only in some cells (Supplementary Fig. 7e, f). Three-dimensional imaging showed that most often, the upper portions of the cell boundary retracted, while lower portions closer to the substrate continued to expand (Supplementary Fig. 7g). Thus, in contrast to the case of single cancer cells dividing in alginate hydrogels, interpolar spindle elongation plays only a minor role in driving mitotic elongation and protrusive force generation.

We next assessed the role of cytokinetic ring contraction in protrusive force generation during mitotic elongation. As cytokinetic ring contraction occurs perpendicular to the mitotic axis, perpendicular inward forces should lead to outward forces along the mitotic axis if volume is conserved or nearly conserved (Fig. 3a). The volume of dividing cells was measured before and after cytokinesis and measured to decrease by only 5% (Fig. 3d). Using computational modelling we evaluated how much a cell can be extended in the axial direction by a contractile force exerted on the equator of the cell (Fig. 3e, f, Supplementary Fig. 8, Supplementary note). Even with a 5% reduction in volume, decreasing ring sizes during cytokinesis were accompanied by 60% elongation along the division axis, suggesting the role of cytokinetic ring contraction in driving mitotic elongation (Fig. 3g). Reductions in volume on the order of 40% would be needed for cytokinetic ring contraction to not drive cell elongation (Fig. 3h). To directly test whether cytokinetic ring contraction is involved in cell elongation, cytokinesis was inhibited by adding blebbistatin ^24^ to cells after they progressed to metaphase, and shortly before mitotic elongation onset (Fig. 3i). Inhibition of cytokinetic ring contraction resulted in binucleate cells and almost completely abrogated cell elongation (60% reduction) and adjacent cell deformation (Fig. 3j, k). Importantly, inhibition of cytokinetic ring contraction also diminished chromosome segregation, highlighting the biological importance of the forces underlying mitotic elongation for proper cell division (Fig. 3l). Similar results were found for inhibition of polo-like kinase 1, which is essential for both anaphase B and cytokinesis progression (Fig. 3m)^25^. Knockdown of anillin within MCF10A cells, resulted in dividing cells contracting their cytokinetic ring to varying degrees, until cytokinesis failure (Supplementary Fig. 9). Knockdown cells exhibited reduced cell elongation during division (Fig. 3n). Furthermore, within knockdown cells, cell elongation was correlated with the extent of maximum cytokinetic ring contraction (Fig. 3o, p). Taken together, these results demonstrate that the forces for mitotic elongation are primarily generated by cytokinetic ring contraction.

### Force generation by mitotic elongation *in vivo*

After identifying mitotic elongation as a direct force generating stage in epithelial monolayers *in vitro,* we examined whether epithelial cells dividing within *in vivo* contexts also exert protrusive forces during mitotic elongation. Consistent deformation of cells adjacent to a mitotic cell during elongation, and an increase in curvature of the cell-cell boundary along the mitotic axis, would indicate protrusive force generation originating from the dividing cell. We first examined early *Drosophila* embryos cell division after the blastoderm stage, focusing on cells within mitotic domains 1 and 5^26^. Dividing cells consistently deformed adjacent rather than perpendicular cells at the center plane, which exhibited reduced cross-sectional areas and increased inward curvatures after division completion (Fig. 4a-d). Next, we examined mitotic events in published literature and analyzed deformation of cell-cell junctions between dividing cells and neighboring cells across a variety of model organisms, including mouse, *Drosophila, Xenopus,* zebrafish, and *C. elegans*^9,26–32^. Negative area strains in adjacent cells were observed in most cases (Fig. 4e). Strikingly, positive changes in curvature of cell-cell boundaries were observed in all instances, with no negative changes in curvature observed, an observation that can only be explained by protrusive force generation from the mitotic cell (Fig. 4f, Supplementary Fig. 10). Interestingly, in the context of stem cells dividing within adult *Drosophila* intestines, the dividing cells recoiled shortly after completing division (Fig. 4g, h). This suggests that the dividing stem cell was bearing compressive forces during mitotic elongation, and upon division completion these forces are dissipated, producing an inward motion of the daughter cells^33^. Thus, the protrusive extracellular forces that drive mitotic elongation appear to be ubiquitous across epithelia.

**Fig. 4.**
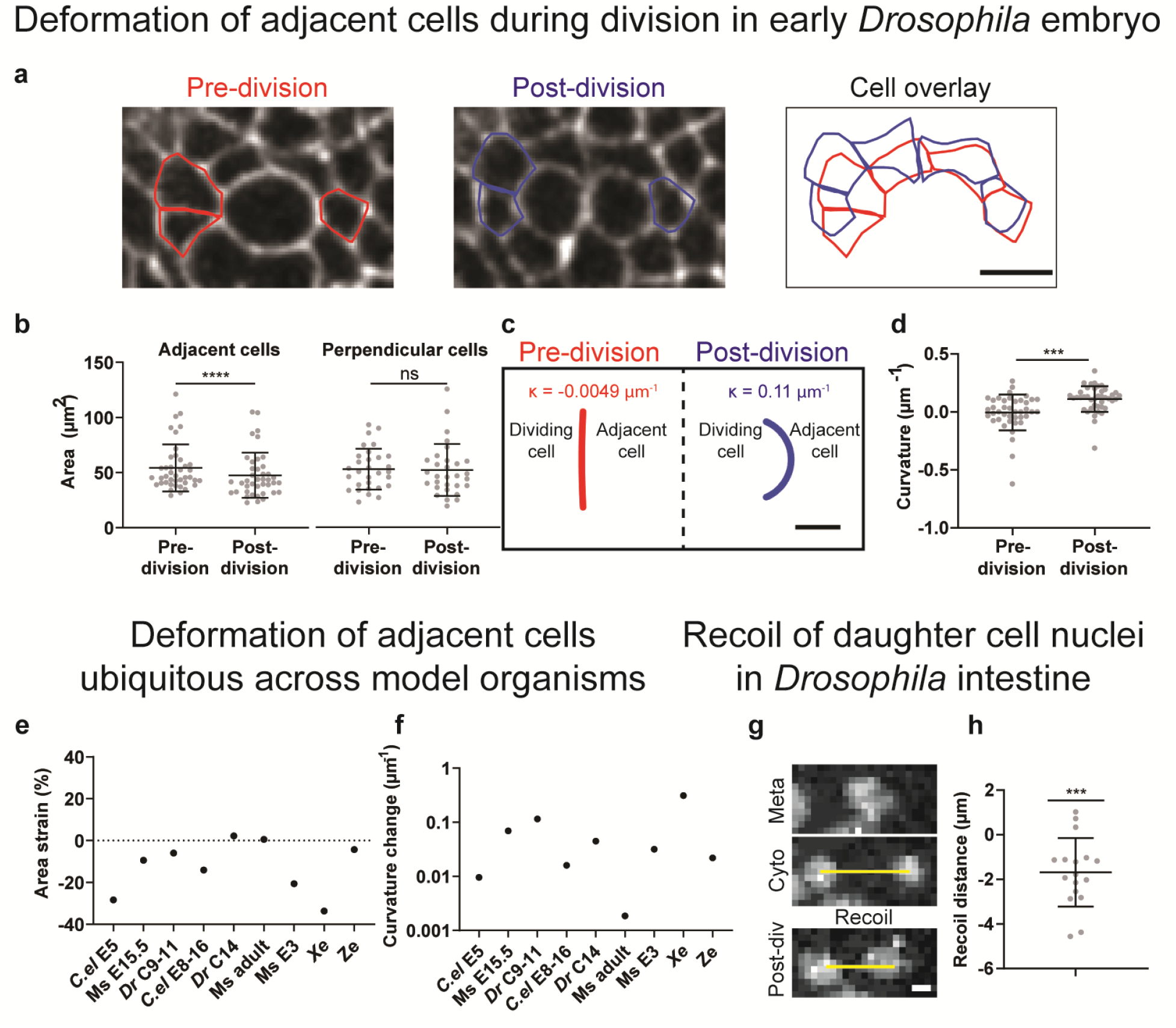
The forces of mitotic elongation are universal across epithelia *in vivo*. **a**, **b**, Deformation of adjacent cells in early *Drosophila* embryos during cell division (**a**), with quantification of adjacent and perpendicular cell area before and after division (**b**). Image in (**a**) is of Utrophin::GFP, which marks the cell cortex. **c**, **d**, Schematic displaying average change in curvature of an adjacent cell during division (**c**) and quantification of curvatures before and after division (**d**). Paired t-tests. Scale bars, 10 μm. **e**, **f**, Deformation of adjacent cells quantified with area strain (**e**) and curvature change (**f**) for *C. elegans* embryo E5 (*C. el*E5), embryonic (E15.5) mouse epidermis (Ms E15.5), *Drosophila* embryo cycles 9-11 (*Dr* C9-11), *C. elegans* embryo E8-16 (*C. el* E8-16), *Drosophila* embryo cycle 14 (*Dr* C14), adult mouse intestinal organoid (Ms adult), embryonic (E3) mouse, *Xenopus* embryo *(Xe),* and zebrafish larva epidermis (Ze). **g**, **h**, Recoil of daughter cell nuclei after division completion within the adult *Drosophila* intestine (**g**), and their quantification (**h**). One-sample t-test. Scale bar, 2.5 μm. P values > 0.05 (n.s.), <0.001 (***), < 0.0001 (****).

## Discussion

Here we showed that cells dividing within epithelial monolayers generate distinct forces in three mitotic stages: mitotic rounding, elongation, and spreading. Inward stresses develop during mitotic rounding, outward stresses are generated along the division axis during mitotic elongation, and uniform outward stresses continue during mitotic spreading. Though the origins of force generation during mitotic rounding, and their implications, have been well studied, mitotic elongation and mitotic spreading have been overlooked. Our results indicate that forces generated during mitotic elongation originate from the dividing cell rather than neighboring cells, with cytokinetic ring contraction playing a greater role in generating protrusive forces than interpolar spindle elongation. This is in contrast to what was observed with single cancer cells dividing within inert hydrogels; where both interpolar spindle elongation and cytokinetic ring contraction were found to play a major role in protrusive force generation^15^. The additional source of protrusive force generation from the interpolar spindle could potentially point to an ability cancer cells must acquire to proliferate in growing tumors that are typically highly compressed and confining^34^. While it has been known that cells have the ability to generate large forces during cell spreading, through the formation of robust adhesions and polymerization of branched actin networks, our work ties these forces to enabling reintegration of daughter cells into the monolayer. Beyond their role in mitosis, mitotic forces have been linked to tissue-scale biological processes such as collective cell migration observed in intestinal crypts and villi^2,5^, cell rearrangements within chick embryos^4^, and invagination within developing *Drosophila* embryos^7,35^. While previous work has typically attributed mitotic forces to mitotic rounding, these processes are inherently anisotropic. Thus, mitotic elongation could play a key, unappreciated role in contributing to, or driving, these important biological processes.

## Materials and Methods

### Cell culture

Parental Madin-Darby canine kidney (MDCK) type II G cells, and MDCK cells stably expressing the nuclear FUCCI cell cycle sensor (gift from Dr. William J. Nelson, Stanford University) or LifeAct::RFP (gift from Dr. Alexander Dunn, Stanford University), were grown in low-glucose Dulbecco’s Modified Eagle Medium (Thermo Fisher, 31600034) supplemented with 10% fetal bovine serum (FBS; GE Healthcare, SH30071.03), 1 g/l sodium bicarbonate, and 100 U/ml penicillin-streptomycin (Thermo Fisher, 15140). Growth media for MDCK cells stably expressing LifeAct::GFP (gift from Dr. Jens Möller, ETH Zurich), E-cadherin::DsRed, truncated E-cadherin (T151-), vinculin::GFP, or α-tubulin::GFP (gifts from Dr. William J. Nelson, Stanford University) was additionally supplemented with 250 μg/ml geneticin (Thermo Fisher, 10131027). T151-cells were cultured with tetracycline-free FBS (GE Healthcare, SH30070.03T)^22^. MCF10A wild-type and LifeAct::RFP mammary epithelial cells (ATCC, CRL-10317) were cultured in DMEM/F12 (Thermo Fisher, 11330057) supplemented with 5% horse serum (Thermo Fisher, 16050122), 20 ng/ml epidermal growth factor (Peprotech, AF-100-15), 0.5 μg/ml hydrocortisone (Sigma H0888), 100 ng/ml cholera toxin (Sigma, C8052), 10 μg/ml insulin (Sigma, 91077C), and 100 U/ml penicillin-streptomycin^36^.

### Polyacrylamide gel fabrication

Polyacrylamide gels were fabricated and functionalized with collagen-I based on a mix of previously published protocols ^16,37–41^. For 1kPa gels (used for all monolayer experiments), a stock solution of 1 ml was made by mixing 100 μl 40% acrylamide (Sigma, A4058), 30 μl 2% bisacrlyamide (Sigma, M1533), 815 μl MilliQ water, and 55 μl of a 1:10 diluted solution of 0.5 μm carboxylate-modified fluorescent beads (Invitrogen, F8888/F8887). Single cells were unable to divide on 1kPa gels and thus were plated on gels previously measured to be 7.43 kPa (250 μl 40% acrylamide, 30 μL 2% bisacrlyamide, 665 μl MilliQ water, and 55 μl beads) ^38^.

### Polyacrylamide gel mechanical characterization

Elastic modulus was measured using an AR-G2 stress-controlled rheometer (TA instruments) equipped with a 25 mm bottom- and upper-plate as described previously ^42^. Polyacrylamide gel solution was deposited on the bottom plate immediately after mixing. A cyclic strain with amplitude 0.01 at a frequency of 1 rad/sec was applied for 1 hr at room temperature (RT) to monitor gel polymerization. The temperature was increased to 37°C and measurements were taken for an additional 15 min. The last 10 recorded values for storage modulus were averaged. This shear modulus value was then converted to an elastic modulus using a poisson’s ratio of 0.457 ^43^.

### Live time-lapse microscopy of cultured cells

Unless stated otherwise, cells were imaged every 1 - 10 min on a Nikon Ti2-E inverted microscope equipped with an ORCA-Flash sCMOS camera, using a 10x or 20x objective, with a pixel size between 0.33 and 0.67 μm. For measuring traction and monolayer stresses for MCF10A cells, cells were imaged every 2 min on a Leica SP8 confocal microscope equipped with a hybrid detector, using a 10x objective, with a pixel size of 0.57 μm. Cells were maintained at 37°C and 5% CO2 for all live-cell experiments. For monolayer height measurements, imaging of cell-matrix adhesions, laser ablation experiments, cell volume measurements, and *Drosophila* experiments, different imaging equipment was used, as described in the respective sections.

### Live time-lapse microscopy of *Drosophila* early embryos

For live imaging of *Drosophila* early embryos, embryos between 3 and 3.5 hours old were dechorionated with 50% bleach, rinsed with water, and mounted, dorsal side up, onto a slide coated with embryo glue (double sided scotch tape dissolved in heptane). No. 1.5 coverslips were used as spacers to create a channel for the mounted embryo. The channel was filled with Halocarbon 27 oil (Sigma). Images were taken of the dorsal side of the head of the embryos when cells in mitotic domains 1 and 5 were dividing. Images were acquired on a Zeiss LSM 710 confocal microscope, with a 40x/1.2 Apochromatic water objective, an argon ion, 561 nm diode, 594 nm HeNe, and 633 nm HeNe lasers. GFP and mCherry markers were simultaneously excited and detected using band-pass filters set at ~490–565 nm for GFP and ~590–690 nm for mCherry. The pinhole was set between 1 and 2 Airy Units for all images. Cell cortices were visualized using a marker for F-actin, the actin binding domain of Utrophin fused to GFP (Utrophin::GFP) ^44^, and microtubules were visualized using tubulin::mCherry.

### Traction force microscopy (TFM)

Cells were removed from gel substrates with 0.5% trypsin-EDTA and reference images of fluorescent beads were taken. Substrate displacements between force and reference gel states were calculated using particle image velocimetry (PIV) implemented with the MATLAB application PIVLab^45^. Two interrogation windows (64 x 64 and 32 x 32 pixels) with 50% overlap were used for most experiments. For MC10A analysis, pixel sizes were greater and thus interrogation windows with sizes 32 x 32 and 16 x 16 were used. Displacements were filtered through a Gaussian filter with standard deviation 6 μm. Substrate displacements were used to calculate cell-matrix stresses using Fourier-transform traction force microscopy derived previously by Trepat et al.^16^, with our results matching that of pyTFM, an opensource TFM tool^46^.

### Monolayer stress microscopy (MSM)

A custom MATLAB script was written to implement MSM, with a monolayer height of 9 μm (Supplementary Fig. 4g, h) as described earlier^17^.

### Analysis of cell-matrix and cell-cell stress data

Dividing cells were manually identified using bright-field or phase imaging. Cell division phases were assigned based on the following standards: start of mitotic rounding–edges of parent cell begin to move inward, metaphase–chromatids were lined along the metaphase plate, late anaphase–chromosomes were moving apart with cytokinetic ring contraction initiated, cytokinesis–cytokinetic ring contraction had completed, and end of post-mitotic spreading–edges of the daughter cells were no longer moving outward. A line was drawn along the long axis of cells at late anaphase to measure the division axis orientation. The corresponding cell-matrix and cell-cell stress data was rotated and centered such that the cell was dividing along the horizontal axis at the center of the map. Data was then fit to a grid with 8.6 μm spacing (11.4 μm for MCF10A analysis). For each dividing cell, cell-matrix and cell-cell stresses were subtracted at two different time points –corresponding to two different cell division stages. Peak traction values were defined as the average of the peak differential stress on the left–hand and right–hand side of the dividing cell, within a 60 μm diameter region of the dividing cell’s center. Center monolayer stress values were defined as the differential stress at the center of the dividing cells. Cell-matrix and cell-cell stress heat maps were generated by averaging the change in stress data from >100 cells.

### Monolayer height measurements

MDCK cells stably expressing LifeAct:RFP were imaged using a laser scanning confocal (Leica, SP8) at varying densities. Volumetric stacks were binarized using FIJI. Height of the monolayer at every point was calculated by taking the difference between the maximum and minimum binarized values.

### Dividing cell edge or vertex categorization

To determine whether cells divide against two adjacent cell edges, two adjacent cell vertices, or an edge and a vertex, phase imaging of MDCK WT or fluorescent imaging of MDCK E-cadherin::DsRed was performed. Each side of the dividing cell was manually categorized as dividing against a cell edge or vertex.

### Truncated E-cadherin experiments

To monitor isolated T151-cells surrounded by MDCK wild-type (WT) cells, mosaic epithelial monolayers were formed using WT and T151-cells at a 95:5 ratio. To minimize the number of adjacent T151-cells formed due to division, an “instant” monolayer was formed as described previously^47^. T151-cells were incubated with 10 μM CellTracker™ Red CMTPX dye (Thermo Fisher, C34552) for 20 min at 37°C prior to trypsinization to differentiate them from MDCK WT cells. After trypsinization, cells were re-suspended in low (5 μM) Ca^2+^ growth media to disrupt cell-cell adhesions and plated at a high (300,000 cells/cm^2^) density in low Ca^2+^ growth media to form an instant monolayer. After 1 hr cell media was replaced with normal growth media (Ca^2+^ 1.8 mM). Imaging began after an additional hour to allow the formation of cell-cell adhesions.

### Truncated E-cadherin immunostaining and imaging

MDCK WT and T151-monolayers were grown on glass substrates conjugated with collagen-I using similar methods outlined in the “Truncated E-cadherin experiments” section. Samples were fixed, permeabilized, blocked, and stained using previously published protocols^48^. Monoclonal rat E-cadherin antibody that specifically binds to the extracellular domain of E-cadherin (Sigma, U3254) was used at 1:1600 dilution, and goat anti-rat Alexa 488- or 555-conjugated secondary antibody (Thermo Fisher, A-11006 and A-21434) was used at 1:1000 dilution, followed by DAPI (Sigma D9542) at 5 μg/ml. Total fluorescence intensity for each image taken was normalized by the maximum fluorescence intensity within a given replicate (either WT or T151-) to account for differences in output between different imaging sessions.

### Neighboring nuclei arrangement, movement, and cross-sectional area change during division

Nuclei were imaged using MDCK cells stably expressing the FUCCI cell-cycle reporter. Nuclei images were binarized and segmented using FIJI. Segmented nuclei were then fit to a grid with 20 μm spacing. Cells within one grid spacing and along the division axis were categorized as adjacent cells. Cells within one grid spacing and along the axis perpendicular to the dividing cell were categorized as perpendicular cells. The arrangement of neighboring cells (one grid spacing away) with respect to the dividing cell’s orientation, was quantified by measuring the angles between the dividing cell at metaphase and neighboring nuclei centroids. Nuclear deformation of cells was quantified by measuring the change in area imaged near the center cross-sectional plane of the cell imaged using wide-field microscopy. Heat maps display average of neighboring cells’ nucleus strain across all observed division events.

### Neighboring cell area measurements during division

For *in vitro* experiments (Fig 2j, k, and n), MDCK cells stably expressing Ecad::DsRed or LifeAct::RFP were imaged. Fluorescent images were binarized and segmented using FIJI. Segmented cells were then fit to a grid with 20 μm spacing. Cells within one grid spacing away from the dividing cell, and along the division axis, were categorized as adjacent cells. Cells within one grid spacing away from the dividing cell, and along the perpendicular axis, were categorized as perpendicular cells. Cells within 100 μm of the dividing cell were categorized as far cells. Deformation of cells was quantified by measuring the change in cross-sectional area imaged near the center plane of the dividing cell using wide-field microscopy. Heat maps display neighboring cells’ area strains averaged across all observed division events. For analysis of cell division within early *Drosophila* embryos after the blastoderm stage (mitotic domains 1 and 5) (Fig. 4a, b), due to the large number of dividing cells within the same region, adjacent or perpendicular cells which had recently divided or were undergoing division soon after, were excluded. Analysis was conducted based on images taken at the center of the dividing cell imaged using laser scanning confocal microscopy. For images displayed from *in vivo* model systems (Fig. 4e), fluorescent images of a single dividing cell with a cell-membrane marker from various model systems were taken from previously published papers ^9,26–32^. Model systems used consist of embryonic (E15.5) mouse epidermis, *Drosophila* embryo (cycles 9 – 11) during segmentation, *Drosophila* embryo after the blastoderm stage (cycle 14), adult mouse intestinal organoid, embryonic (E3) mouse, *Xenopus* embryo during neural tube closure, *C. elegans* embryo (E5), *C. elegans* embryo intestine (E8-E16), and zebrafish larva epidermis. Images were analyzed similarly but with manual outlining. Area strains displayed are from a single division event and average measurements from 2 to 4 adjacent cells. For the analysis of zebrafish larvae epidermis, image scale was estimated based on another image of zebrafish ^28^.

### Curvature (κ) measurements of cell-cell boundaries during division

Common boundary points between the dividing cell and neighboring cells were noted. If the angle between the division axis and the axis defined by a daughter cell’s centroid and the centroid of the common boundary points was below 100°, the neighboring cell was considered to be an adjacent cell. Common points between the dividing cell and an adjacent cell were fit to a circle using code from the MATLAB central file exchange utilizing least-squares fitting ^49,50^. The inverse of the radius of the fitted circle was taken to be the curvature value. If the center of the fitted circle was not in the direction of the dividing cell, the curvature was taken to be negative. Representative arcs were drawn at arbitrary spacing, with arbitrary, but equal lengths between the two time points compared, and with average curvature values of measured data.

### Adhesion imaging and tracking

To image and track adhesions, total internal reflection microscopy (TIRF) was performed using a Nikon spinning disk confocal microscope at the Stanford Cell Sciences Imaging Facility. Adhesions of dividing and neighboring cells were imaged using MDCK cells stably expressing vinculin::GFP. For each dividing cell, adhesions within a 50 x 30 μm (long axis oriented along division axis) perimeter were tracked as the dividing cell transitions from metaphase to cytokinesis completion. The open-source Focal Adhesion Analysis Server was used to identify adhesions and measure their displacement^51^. We measured the number of adhesions identified when the dividing cell reaches metaphase and cytokinesis completion, as well as how far adhesions were moving away or toward the dividing cell along the division axis between these two division stages.

### Pharmacological inhibition experiments to assess the role of interpolar spindle elongation and cytokinetic ring contraction

Cell elongation and interpolar spindle elongation during division was measured by imaging MDCK cells stably expressing either E-cadherin::DsRed or LifeAct::RFP, and α-tubulin::GFP, respectively. Chromosome separation was visualized using brightfield imaging. Blebbistatin (100 μM, Sigma, 203389), BRD9876 (40 μM, Tocris, 9876), and BTO-1 (50 μM, Sigma, B6311) were used to inhibit myosin II, kinesin-5, and polo-like kinase 1, respectively^23–25^. To specifically evaluate the contribution of interpolar spindle elongation and cytokinetic ring contraction during mitotic elongation, only cells that had progressed to metaphase after the drug was added were considered. Only fields of view with similar monolayer densities were compared between control and experimental conditions. For assessing the role of interpolar spindle elongation, cell elongation was measured 6 min after anaphase onset, and for assessing the role of cytokinesis, cell elongation was measured 8 – 10 min after anaphase onset. For cells treated with blebbistatin or BTO-1, only cells that failed to complete cytokinesis and had become binucleate were considered. Reduction in cell elongation was defined as the percent difference in mean cell elongation from experimental and control conditions, normalized to the control condition.

### Anillin knockdown

Knockdown of anillin (ANLN) within MCF10A cells stably expressing LifeAct::RFP was accomplished using Dharmacon ON-TARGETplus Human ANLN siRNA SmartPool (LQ-006838-00-0002). A non-targeting siRNA was used as a control (CTRL). Cells were transfected with a final siRNA concentration of 20 nM using Dharmacon 1 reagent. Cells were imaged and assayed for Western blot 24 to 48 hours after transfection. Cell division within both control and experimental groups was slightly delayed, and thus cell elongation was measured 12 min after anaphase onset. Within the siANLN experimental group, only dividing cells that resulted in cytokinesis failure were considered. For correlation between cytokinetic ring contraction and cell elongation within siANLN dividing cells, minimum cell width was measured along the cytokinetic ring, and cell elongation was measured at the time cytokinetic ring contraction reached a minimum.

### Western blot of anillin knockdown

Cells transfected with siCTRL or siANLN were lysed and denatured, followed by gel electrophoresis, transfer to nitrocellulose, blocking, primary antibody incubation, and secondary antibody incubation following previous protocols^48^. 50 μg of total protein was loaded per lane, primary anillin antibody was used at 200 ng/ml (Bethyl Laboratories, A301-405A), and secondary antibody (Li-Cor Biosciences, 926-32213) at 1:10,000 dilution. Protein concentration was compared by normalizing experimental band intensity to control intensity (relative density), and then normalizing the anillin relative density to that of a loading control (GAPDH; abcam, 181602), which was imaged at a lower intensity to avoid saturation.

### Laser ablation

Laser ablation was performed on a laser scanning confocal microscope (Zeiss LSM 780) at the Stanford Cell Sciences Imaging Facility using a Mai Tai DeepSee (Spectra-Physics) multi-photon laser (800 nm wavelength). MDCK cells stably expressing α-tubulin::GFP were maintained in cell-culture conditions at 37°C with 5% CO2. After identifying dividing cells in late anaphase, a region of interest (ROI) at the center of the dividing cell was drawn. Ablation was performed by scanning the multi-photon laser within the ROI at 5 – 7 z-planes (1 – 2 μm apart), spanning the thickness of the dividing cell. Before and after ablation images were taken at the center plane. Change in distance between the dividing cell’s chromosomes and length were measured. For 3D membrane imaging, cells were stained with CellMask™ orange plasma membrane stain (Thermo Fisher, C10045) at 5μg/ml for 10 min prior to imaging. Before and after ablation z-stacks were taken with 0.75 – 1 μm spacing between z-slices. Images displayed are cross-sectional images of the dividing cell’s center (Supplementary Fig. 7g). The boundary of the dividing cell at pre- and post-ablation time points was outlined for comparison.

### Cell volume measurements

MDCK cells stably expressing LifeAct::GFP or LifeAct::RFP were imaged using a laser scanning confocal (Leica, SP8). Cells at metaphase were identified using a nuclear stain (Thermo Fisher, R37106). Volumetric stacks were taken immediately after anaphase onset, at the start of cytokinesis, and upon cytokinesis completion. Cell volume was measured by calculating the number of voxels within the cell region by manually outlining the dividing cell at each imaging plane.

### Computational modeling of cell elongation during cytokinesis

Using a computational model, we evaluated how much a cell can be elongated in the axial direction due to a contractile force exerted on the equator of the cell. The details of the computational model are explained in the Supplementary note. The cell is simplified into a three-dimensional structure consisting of a membrane with the conservation of volume and area. The membrane is coarse-grained using a triangulated mesh as in previous works ^52^. Extensional stiffness prevents chains between nodes on the mesh from elongating or shortening to a very large extent. Bending stiffness maintains a dihedral angle formed by adjacent triangles on the mesh near equilibrium level. Volume encapsulated by the membrane and the total surface area of the membrane are maintained near their initial values to various extents. The membrane initially has a spherical shape whose radius is 10 μm. To mimic the constriction of a cytokinetic ring, membrane nodes located near the equator are displaced at constant speed (Fig. 3f). Consequently, the narrow region contracts toward the cytokinetic axis as the constricting ring observed during cytokinesis. We ran simulations with different strengths of volume and area conservation and measured how much the cell-like structure is elongated in the axial direction.

### Calculation of nuclei recoil distance in adult *Drosophila* intestine

Images of nuclei within the intestine of adult *Drosophila* were taken from previously published data ^33^. The nuclei of dividing mother stem cell and resulting daughter stem cells were segmented using Imaris. The recoil distance was calculated as the difference in distances of daughter cell’s nuclei centroids, between division completion, and 7.5 min later.

### Statistics

All statistical analysis and graphical figures were done using GraphPad Prism or MATLAB. All statistical tests used and information on replicates is summarized in Table S1. All data points within different replicates were combined before statistical tests were performed. For graphs with error bars, the center bar indicates mean and upper and lower bars indicate standard deviation.

## Data and code availability

The main data supporting the results presented are available within the paper and all other data and code are available from the corresponding author upon request.

## Acknowledgements

The authors would like to acknowledge help from members of the Chaudhuri Laboratory, and Dr. Marc Levenston (Stanford University) for use of the rheometer. The authors thank Dr. Xavier Trepat and Dr. Manuel Gómez González (Institute of Bioengineering of Catalonia) for assistance with traction force microscopy code. The authors would like to thank the Stanford Cell Sciences Imaging Facility for use of Imaris imaging software, Zeiss laser scanning microscope 780, and Nikon spinning disk confocal microscope.

## Author contributions

O.C. and V.K.G. conceived the study, designed experiments, and wrote the manuscript. V.K.G. performed most experiments and data analysis. S.N. assisted with gel mechanical characterization, laser ablation experiments, TFM calculations, and MSM calculations. T.K. performed simulations of volume change during cytokinesis. A.C.M. and J.C. provided and assisted with the analysis of early *Drosophila* embryo videos. L.E.O.B., J.L.M., and E.N.S provided and assisted with the analysis of adult *Drosophila* intestine videos.

