## Supplementary Material for "The nature of mitotic forces in epithelial monolayers"

**This PDF file includes:**

Supplementary note
Supplementary figures 1 - 10
Supplementary tables 1 – 2
Supplemental references

.

**Supplementary note for computational modeling of cell elongation during cytokinesis**

In the model, cell division is simulated using a three-dimensional structure comprised of an initially spherical membrane with the conservation of volume and area. The membrane is simplified into a triangulated mesh as in previous works [1]. The number of nodes consisting the membrane mesh is 10,242, and the initial length of chains between adjacent nodes is ~0.5 μm.

**Brownian dynamics with the Langevin equation**

Displacements of membrane nodes are determined by the Langevin equation with inertia neglected:

$$\mathbf{F}_i - \zeta_i \frac{\mathrm{d}\mathbf{r}_i}{\mathrm{d}t} + \mathbf{F}_i^{\mathrm{T}} = 0 \qquad \text{(S1)}$$

where $\mathbf{r}_i$ is a position vector of the $i$th node, $\zeta_i$ is a drag coefficient, $t$ is time, $\mathbf{F}_i$ is a deterministic force, and $\mathbf{F}_i^{\mathrm{T}}$ is a stochastic force satisfying the fluctuation-dissipation theorem [2]:

$$\left\langle \mathbf{F}_i^{\mathbf{T}}(t)\mathbf{F}_j^{\mathbf{T}}(t) \right\rangle = \frac{2k_{\mathbf{B}}T\zeta_i\delta_{ij}}{\Delta t}\boldsymbol{\delta} \qquad \text{(S2)}$$

where $\delta_{ij}$ is the Kronecker delta, $\boldsymbol{\delta}$ is a second-order tensor, and $\Delta t$ is a time step.

The drag coefficients of membrane nodes are calculated as follows:

$$\zeta_i = \lambda A_i \qquad \text{(S3)}$$

where $\lambda$ is a constant, and $A_{\mathrm{i}}$ is the instantaneous area of a pentagon or a hexagon whose center corresponds to a node $i$ (Supplementary Fig. 8a).

Positions of all nodes are updated via Euler integration scheme:

$$\mathbf{r}_i(t+\Delta t) = \mathbf{r}_i(t) + \frac{\mathrm{d}\mathbf{r}_i}{\mathrm{d}t}\Delta t = \mathbf{r}_i(t) + \frac{1}{\zeta_i}\left(\mathbf{F}_i + \mathbf{F}_i^{\mathrm{T}}\right)\Delta t \qquad \text{(S4)}$$

**Extensional and bending forces**

Extensional stiffness ($\kappa_{\mathrm{s}}$) keeps chain length between nodes remaining within a given range; it is allowed to vary between 0.05 μm and 1.25 μm without resistance, but if the chain length increases or decreases beyond the range, a linear spring force is applied to bring it back within the range. Thus, the potential function ($U_{\mathrm{s}}$) for the chain extension is:

$$U_{\mathrm{s}} = \begin{cases} \frac{1}{2}\kappa_{\mathrm{s}}\left(r - r_{0,\mathrm{L}}\right)^2 & \text{if } r < r_{0,\mathrm{L}} \\ 0 & \text{if } r_{0,\mathrm{L}} \le r \le r_{0,\mathrm{H}} \\ \frac{1}{2}\kappa_{\mathrm{s}}\left(r - r_{0,\mathrm{H}}\right)^2 & \text{if } r > r_{0,\mathrm{H}} \end{cases} \qquad \text{(S5)}$$

where $r$ is instantaneous chain length, and $r_{0,\mathrm{L}}$ and $r_{0,\mathrm{H}}$ are the lower and upper limits of the range.

Bending stiffness ($\kappa_{\mathrm{b}}$) maintains a dihedral angle ($\theta$) formed by two adjacent triangles located on the mesh near equilibrium level ($\theta_0$ = 0 rad):

$$U_{\mathrm{b}} = \kappa_{\mathrm{b}}\left[1 - \cos\left(\theta - \theta_0\right)\right] \qquad \text{(S6)}$$

Forces calculated from $U_{\mathrm{b}}$ are applied to four nodes that constitute the two adjacent triangles.

**Conservation of volume and surface area**

Volume encapsulated by the membrane is conserved by the following potential ($U_{\mathrm{v}}$):

$$U_{\mathrm{v}} = \frac{\kappa_{\mathrm{v}}\left(V - V_0\right)^2}{2V_0} \tag{S7}$$

where $\kappa_{\mathrm{v}}$ is the strength of volume conservation, and $V$ and $V_0$ are instantaneous and equilibrium volume within the membrane. The global volume is calculated by summing the volume of tetrahedra, each of which is defined by three nodes of a triangle on the mesh and the center point of the membrane. For maintaining $V$ near $V_0$, forces calculated from $U_{\mathrm{v}}$ move triangles on the mesh outward or inward in directions normal to the triangles. For example, if $V$ is smaller than $V_0$, forces are applied to membrane nodes to push triangles on the mesh outward so that $V$ can increase.

The global surface area of the membrane is conserved by another potential ($U_{\mathrm{a}}$):

$$U_{\mathrm{a}} = \frac{\kappa_{\mathrm{a}}\left(A - A_0\right)^2}{2A_0} \tag{S8}$$

where $\kappa_{\mathrm{a}}$ is the strength of area conservation, and $A$ and $A_0$ are instantaneous and equilibrium surface area. Forces calculated from $U_{\mathrm{a}}$ make individual triangles larger or smaller to maintain $A$ near $A_0$. For example, if $A$ is smaller than $A_0$, forces are applied to membrane nodes to make triangles larger so that $A$ can increase. It has been shown that the surface area of cells can be increased due to membrane reservoirs [3,4], whereas lipid vesicles commonly used for in vitro experiments cannot accommodate a large change in the surface area due to the absence of membrane reservoirs.

**Simulation setup**

In each simulation, a spherical membrane whose radius is 10 μm is located at the center of a large three-dimensional domain (100×100×100 μm). To mimic the constriction of a cytokinetic ring, 522 membrane nodes within a region whose width is 1 μm near the equator of the membrane are displaced inward toward the cytokinetic axis at a constant speed, $v_{\mathrm{c}}$ (Fig. 3f). Those membrane nodes stop being displaced after a distance to the cytokinetic axis becomes less than 0.5 μm. Consequently, the narrow region contracts toward the cytokinetic axis as the constricting ring observed during cytokinesis.

We ran simulations with different strengths of volume and area conservation and evaluated how much the cell-like structure is elongated in the axial direction. The extent of cell elongation is defined by dividing the distance between the leftmost and rightmost points of the membrane by the initial diameter of the membrane.

**Supplementary Fig. 1**

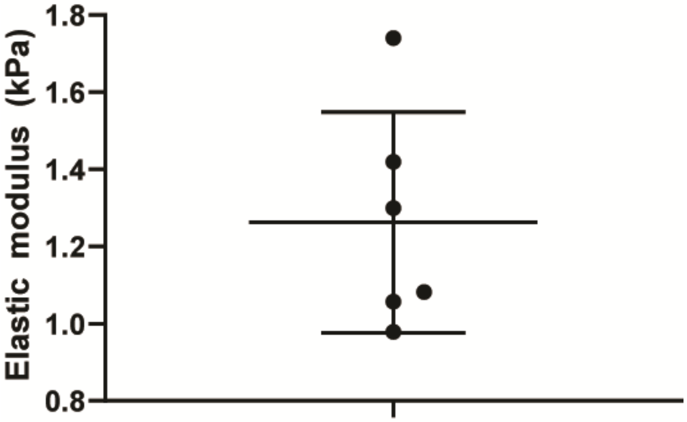


**Mechanical characterization of polyacrylamide gels.** Measurements of elastic moduli of 6 independent polyacrylamide gel samples fabricated from the same formulation.

### Supplementary Fig. 2

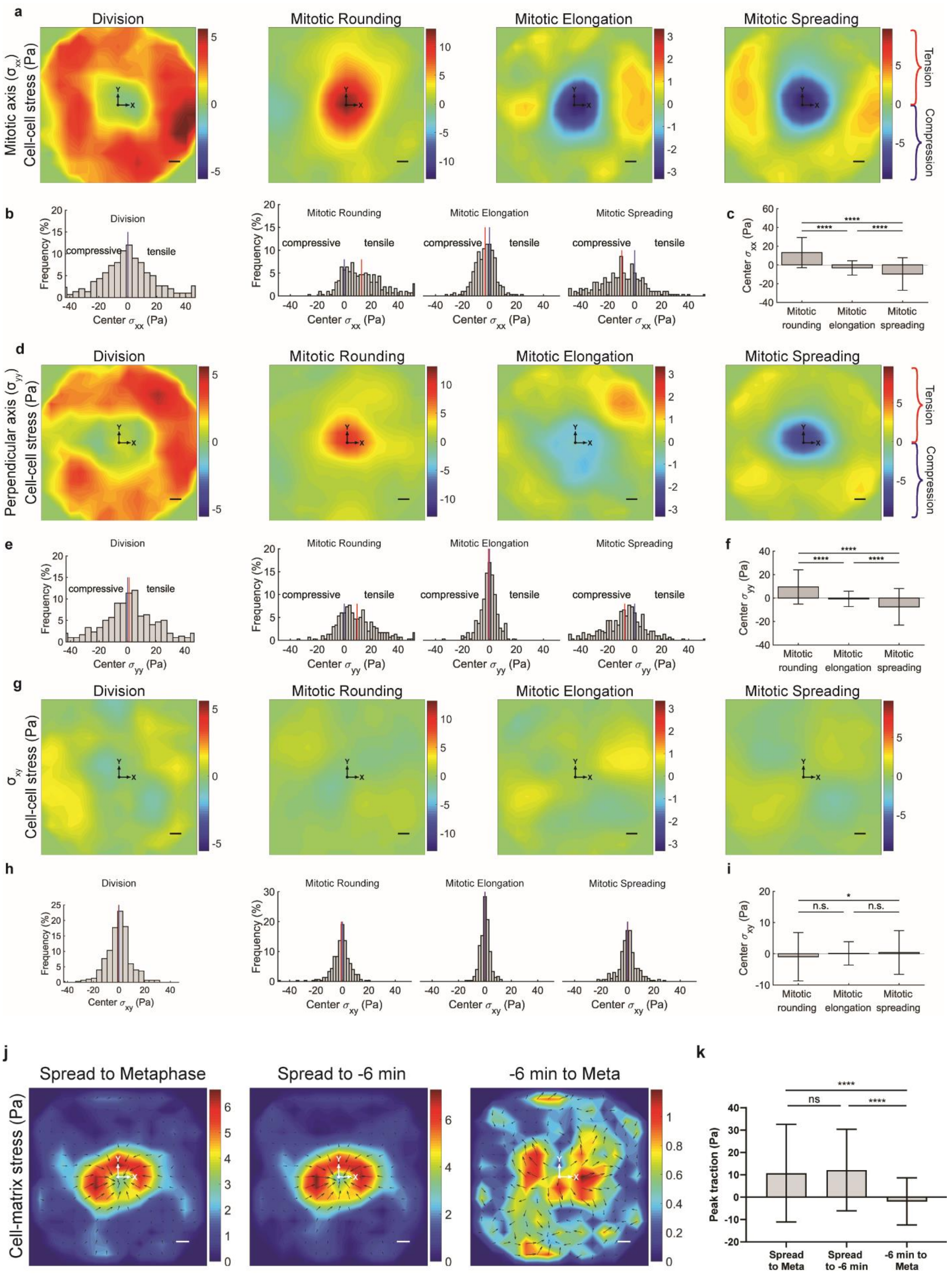

**MDCK epithelial division is accompanied by distinct stages of cell-cell stress. a-i,** Change in cell-cell stress along division axis (**a**-**c**), perpendicular axis (**d**-**f**), and shear directions (**g**-**i**) during division represented as averaged heat maps (**a**, **d**, **g**), distributions (**b**, **e**, **f**), and average-value comparisons (**c**, **f**, **i**) for entire division process, mitotic rounding, mitotic elongation, and mitotic spreading. Mitotic rounding consists of inward and outward force generation stages. **j**, **k**, Average change in cell-matrix stress during the course of mitotic rounding (from a spread parent cell to metaphase), from a spread parent cell to 8 min prior to metaphase, and from 8 min prior to metaphase to metaphase (**j**), and corresponding quantification (**k**). For comparisons, Multiple comparison test P values > 0.05 (n.s.), <0.05 (*), <0.001 (***), <0.0001 (****). Scale bars, 10 µm.

**Supplementary Fig. 3**

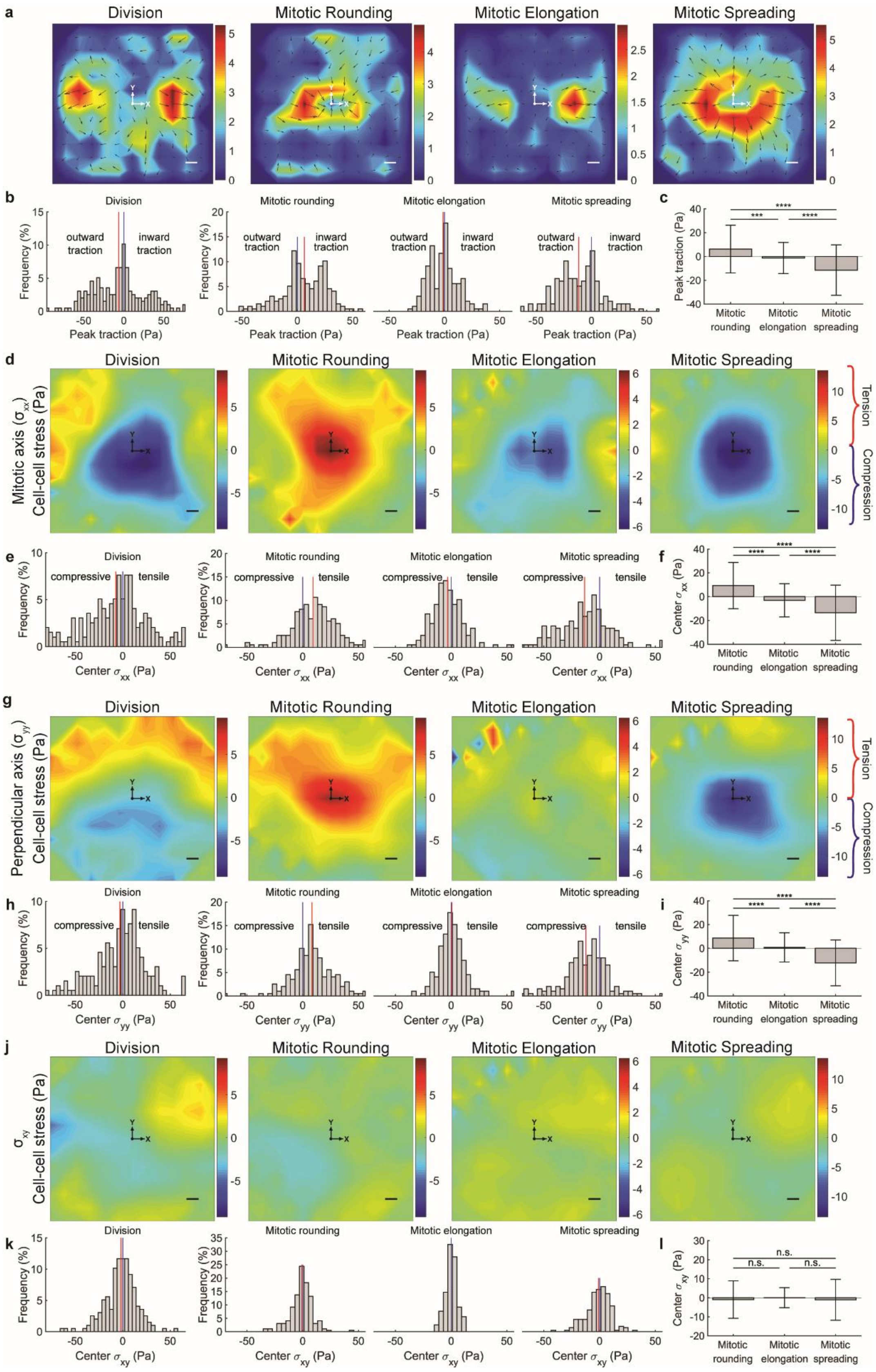

**MCF10A epithelial division is accompanied by distinct stages of cell-matrix and cell-cell stress. a-l**, Change in cell-matrix stress (**a-c**) and cell-cell stress during division along division axis (**d-f**), perpendicular axis (**g-i**), and shear directions (**j-l**) represented as averaged heat maps (**a**, **d**, **g**, **i**), distributions (**b**, **e**, **h**, **k**), and average-value comparisons (**c**, **f**, **i**, **l**) for entire division process, mitotic rounding, mitotic elongation, and mitotic spreading. For comparisons, Multiple comparison test P values > 0.05 (n.s.), , <0.01 (**), <0.001 (***), <0.0001 (****). Scale bars, 10 µm.

**Supplementary Fig. 4**

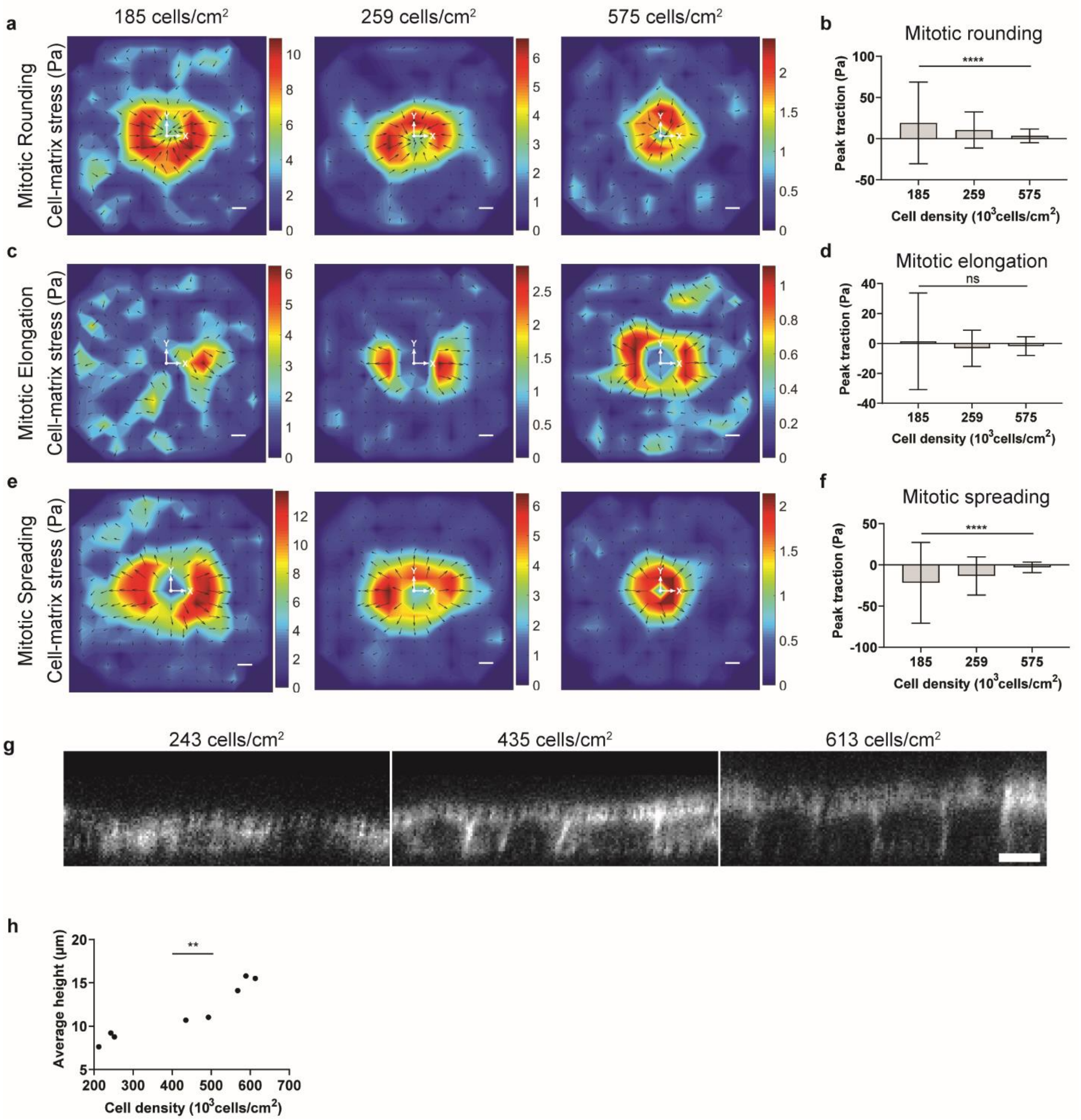


**Increasing epithelial monolayer densities are correlated with reduced measured mitotic forces and increasing heights. a-f**, Change in cell-matrix stress during division at varying MDCK monolayer densities represented as averaged heat maps and average-value comparisons, respectively, for mitotic rounding (**a-b**), mitotic elongation (**c-d**), and mitotic spreading (**e-f**). One-way ANOVA post-test for trend P values > 0.05 (n.s.), <0.0001 (****). **g-h**, Representative cross-sectional images of MDCK cells stably expressing LifeAct:RFP at varying densities (**g**) and their height quantification (**h**). Spearman's rank P value < 0.01 (*). The increased cell heights and reduced cross-sectional areas at greater monolayer densities make the plane stress assumption used in monolayer stress microscopy (MSM) less valid, which could explain why measured mitotic stresses decrease at increasing monolayer densities [5]. Scale bars, 10 µm.

**Supplementary Fig. 5**

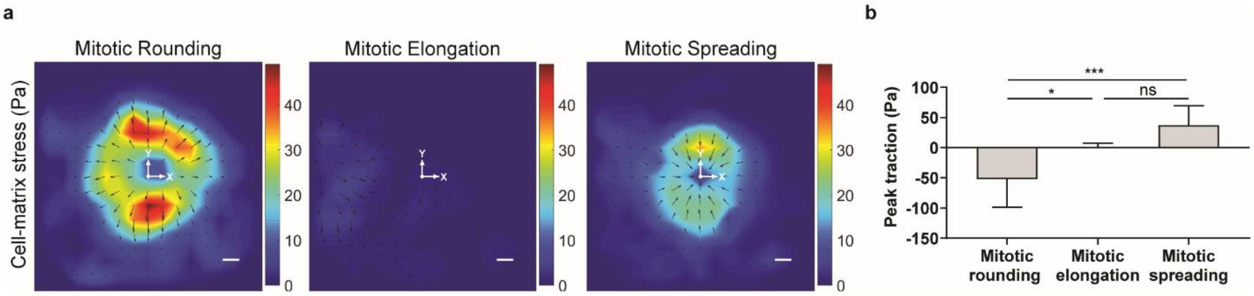


**Single-cell division is accompanied by distinct stages of cell-matrix stress. a-b**, Average change in cell-matrix stress during mitotic rounding, mitotic elongation, and mitotic spreading (**a**), and their quantification (**b**). Compared to division within epithelial monolayers, during single-cell division, cell-matrix stresses are directed outward during mitotic rounding and inward during mitotic spreading, with no discernable stress pattern during mitotic elongation. Scale bar, 10 µm. Tukey's multiple comparison test. P values > 0.05 (n.s.), < 0.05 (*), <0.001 (***).

**Supplementary Fig. 6**

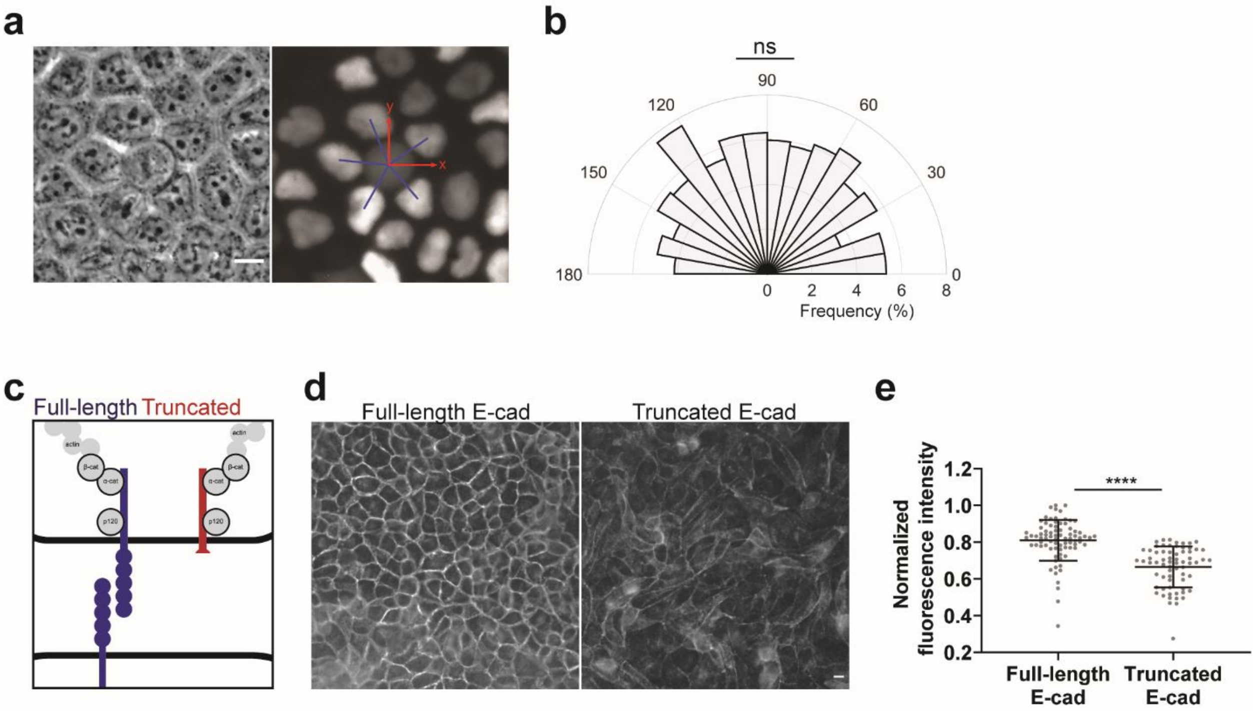


**Adjacent cells do not pull on dividing cells.** Arrangement of neighboring nuclei during division is random. **a**, Representative brightfield and fluorescent image of MDCK cells stably expressing the nuclear FUCCI cell-cycle reporter, with dividing cell centered, and blue lines indicating angles between dividing cell and neighboring cells. **b**, Distribution of angles between dividing and neighboring cells. $\chi^2$ test against difference from uniform distribution. **c**, Cartoon depicting structure of full-length and truncated E-cadherin protein [6]. **d**, **e**, MDCK cells with dominant expression of truncated E-cadherin exhibit reduced levels of extracellular E-cadherin. Representative images of fluorescent stain of extracellular E-cadherin for MDCK wild-type cells (full-length) and MDCK cells stably expressing truncated E-cadherin (**d**) and the fluorescent intensity quantification (**e**). Unpaired t-test. P values > 0.05 (n.s.), < 0.0001(****). Scale bars, 10 µm.

**Supplementary Fig. 7**

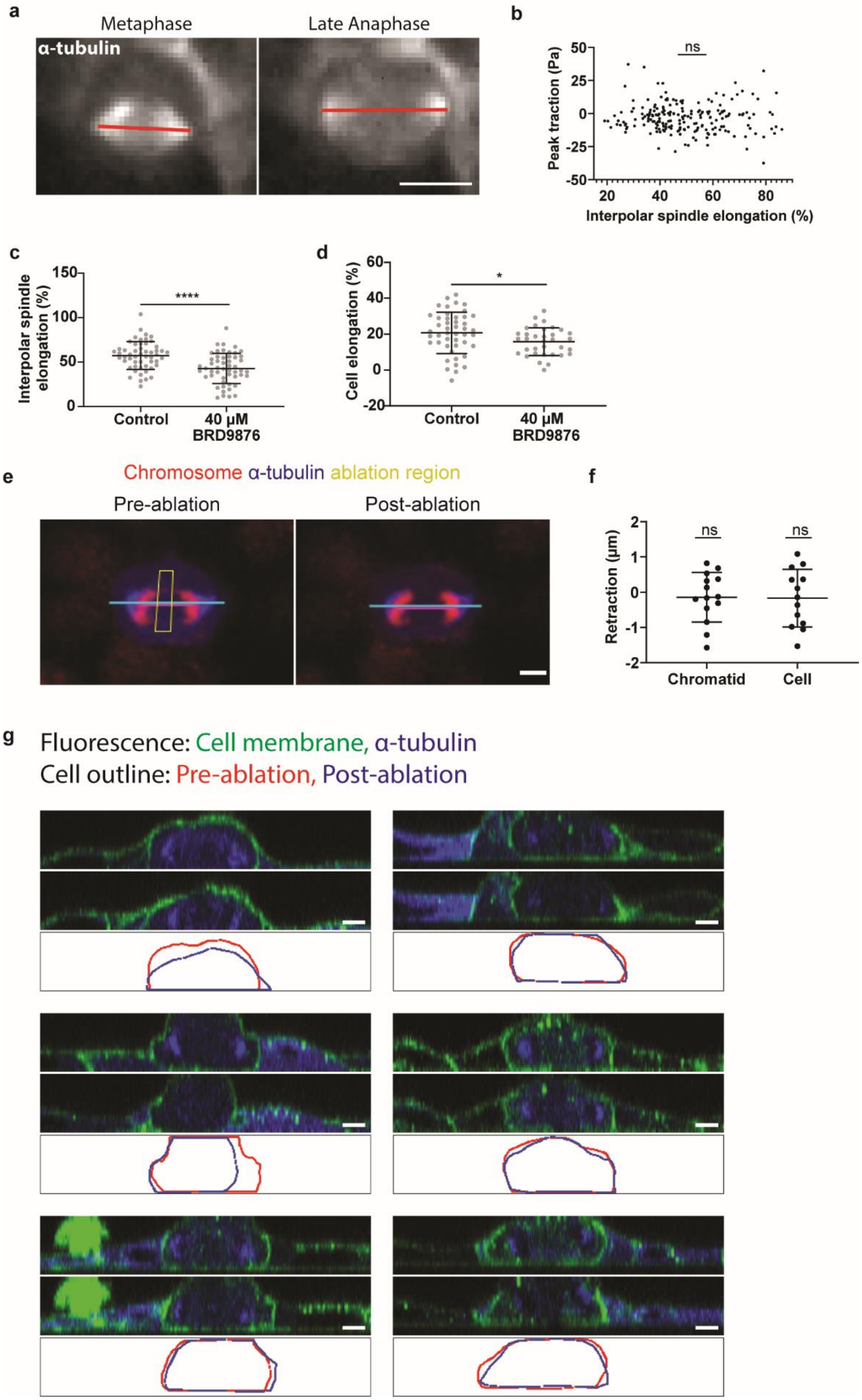

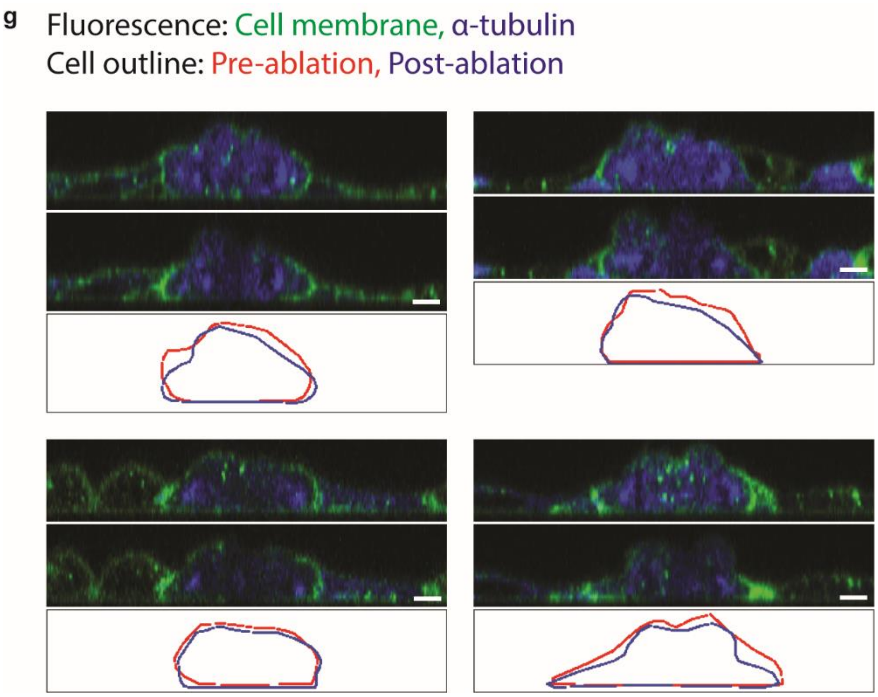


**Forces for mitotic elongations are not generated primarily from interpolar spindle elongation. a**, Quantification of interpolar spindle elongation between metaphase and late anaphase. Scale bar, 10 µm. **b**, Correlation between interpolar spindle elongation and peak traction stress from metaphase to late anaphase. Pearson's *r*. **c**, **d**, Interpolar spindle elongation (**c**) and cell elongation quantification (**d**) with and without inhibition of kinesin-5 with BRD9876 treatment. Unpaired t-tests. **e**, Laser ablation reveals the interpolar spindle does not consistently bear compression. Representative image of dividing MDCK cell before and after ablation of the interpolar spindle, with magenta line indicating distance between sister chromosomes and cyan line indicating cell length. Scale bar, 5 µm. **f**, Quantification of change in length between sister chromosomes and the dividing cell before and after ablation. One-sample t-test. **g**, Three-dimensional imaging of ablated interpolar spindle results in retraction in only upper regions of the dividing cell. Array of cross-sectional images of MDCK cells before and after ablation. Scale bars, 5 µm. P values > 0.05 (n.s.), < 0.05 (*), <0.0001 (****).

**Supplementary Fig. 8**

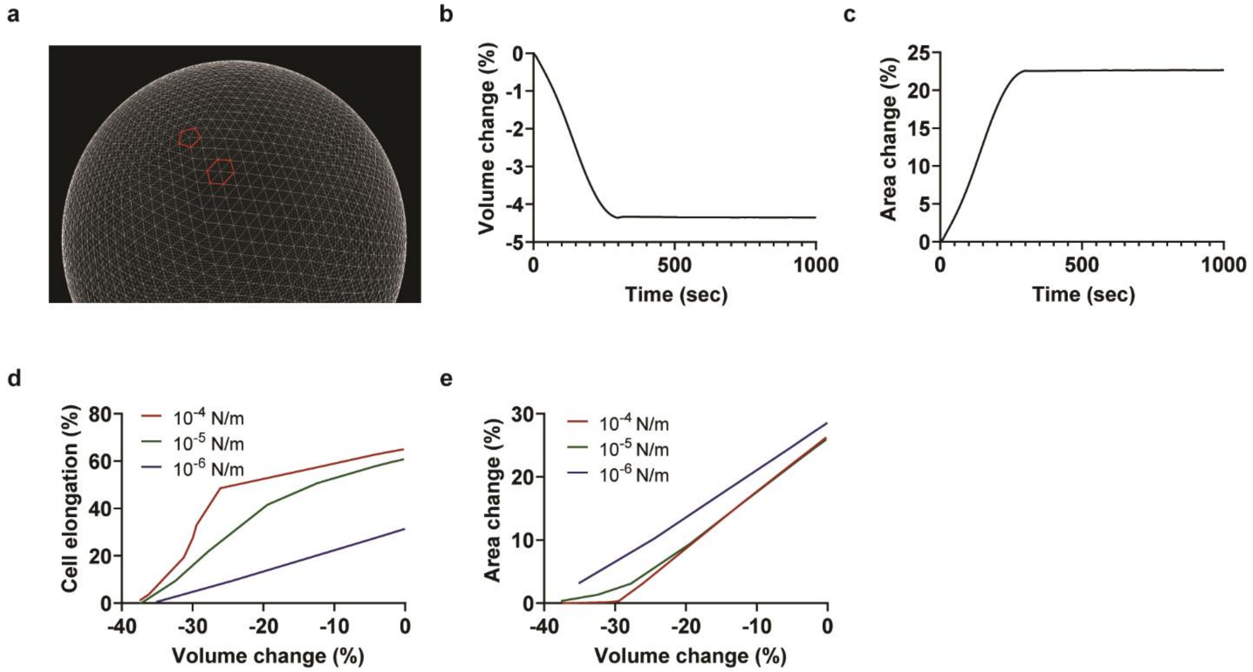


**Simulations reveal axial elongation due to varying levels of volume and area conservation during division. a**, A snapshot depicting coarse-grained triangular mesh consisting of pentagons and hexagons. **b**, **c**, Volume change (**b**) and area change (**c**) during the progression of cytokinetic ring contraction corresponding to the case shown in Fig. 3, e to g. **d**, **e**, Final cell elongation vs final volume change (**d**) and final area change vs final volume change (**e**) for varying degrees of area conservation.

**Supplementary Fig. 9**

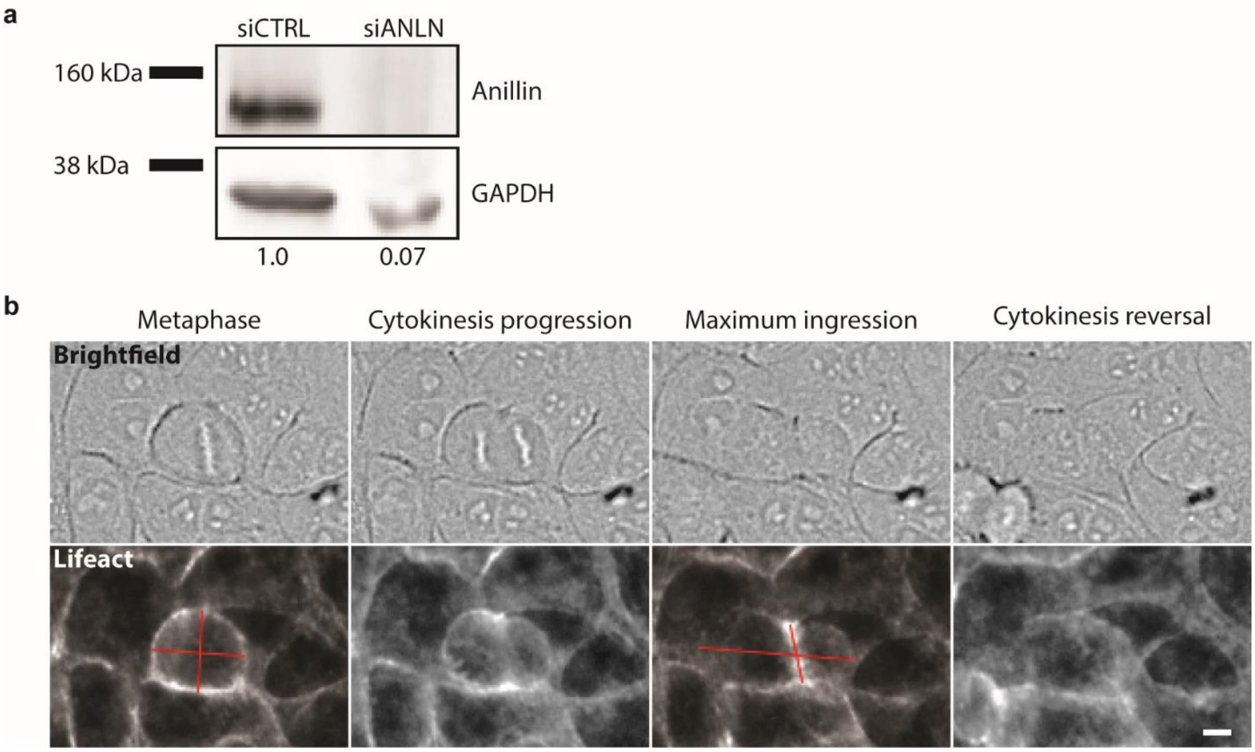


**Knockdown of anillin within MCF10A cells. a**, Western blot indicating knockdown of anillin. Anillin and GAPDH were imaged at different intensities. **b**, Division time-course displaying cell treated with siRNA anillin fails to complete cytokinesis. Scale bar, 10 µm.

**Supplementary Fig. 10**

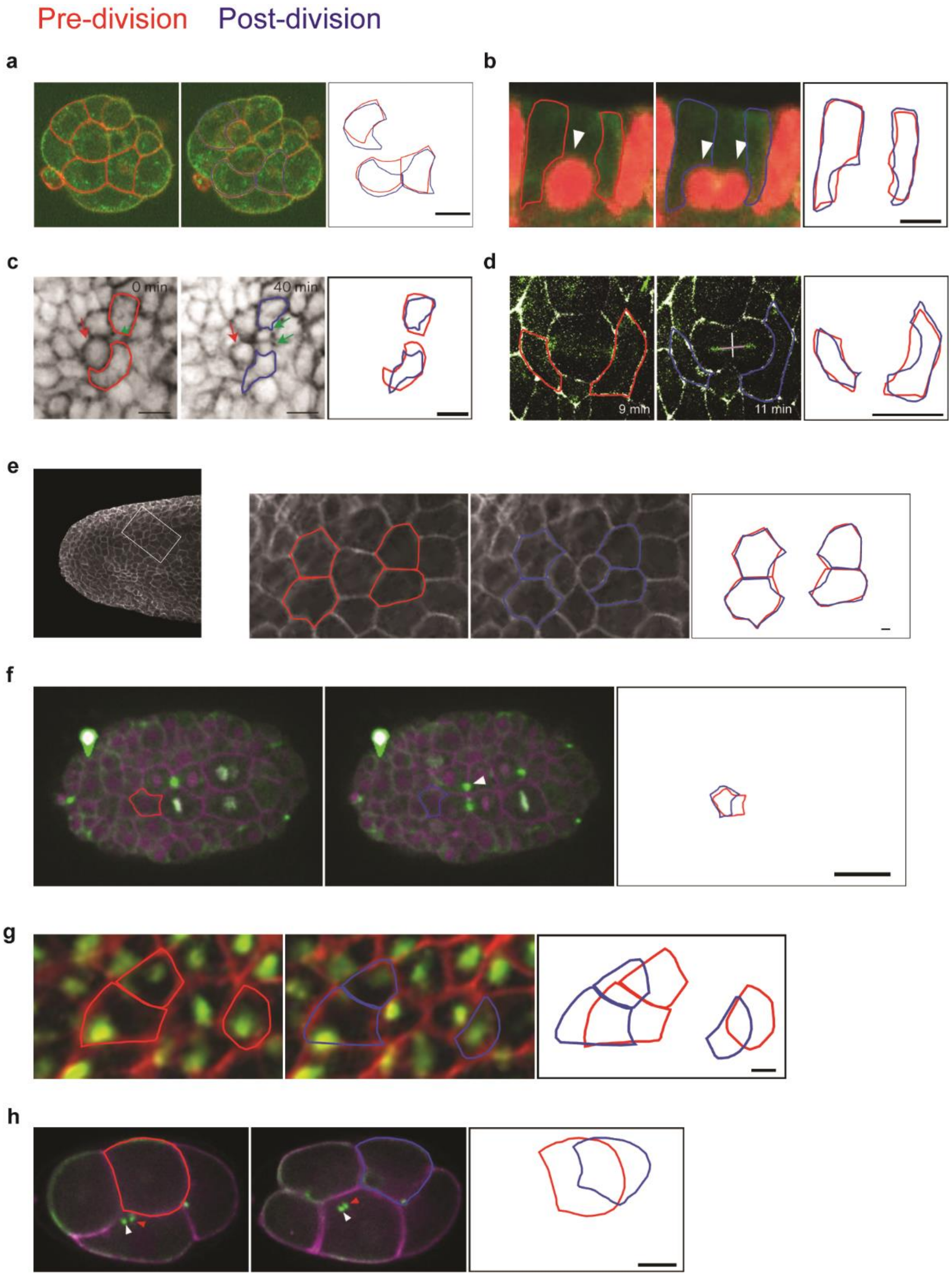


**The forces of mitotic elongation are universal across epithelia *in vivo*. a-h**, Images of adjacent cell deformation during division within mouse blastocyst (E3) (**a**), adult mouse intestinal organoid (**b**), embryonic (E15.5) mouse epidermis (**c**), *Drosophila* embryo cycles 9-11

(**d**), zebrafish larva epidermis (**e**), *C. elegans* embryo E8-16 (**f**), *Xenopus* embryo (**g**), and *C. elegans* embryo E5 (**h**). Scale bars, 15 µm (**c**), 10 µm (**a**-**b**, **d**-**h**).

**Supplementary table 1.**
Information on replicates, number of data points, and statistical testing for all data shown. N/A: not applicable.

| Figure | | Experiment | Biological Replicates | Number of cells | Statistical test |
|---|---|---|---|---|---|
| **1** | **a** | Cartoon | N/A | N/A | N/A |
| | **b** | Cartoon | N/A | N/A | N/A |
| | **c** | Representative image | N/A | N/A | N/A |
| | **d** | Heat map | 3 | 300 dividing cells | N/A |
| | **e** | Representative image | N/A | N/A | N/A |
| | **f** | Heat map | 3 | 300 dividing cells | N/A |
| | **g** | Frequency distribution | 3 | 300 dividing cells | None |
| | **h** | Distribution comparison | 3 | 300 dividing cells | Multiple comparison test |
| **2** | **a** | Cartoon | N/A | N/A | N/A |
| | **b** | Pie chart | 3 | 212 dividing cells | None |
| | **c** | Heat map | 2 | 116 dividing cells in each experimental group | N/A |
| | **d** | Distribution comparison | 2 | 116 dividing cells in each experimental group | Two-sample t-test |
| | **e** | Representative image | N/A | N/A | N/A |
| | **f** | Frequency distribution | 3 | 152 dividing cells; 261 adjacent nuclei | One-sample t-test |
| | **g** | Distribution comparison | 4 | 12 cells | Paired sample t-test |
| | **h** | Representative image | N/A | N/A | N/A |
| | **i** | Frequency distribution | 4 | 12 dividing cells; 2724 tracked adhesions | One-sample t-test |
| | **j** | Representative image | N/A | N/A | N/A |
| | **k** | Heat map | 2 | 200 dividing cells; 369 adjacent cells; 369 perpendicular cells; 12,371 neighboring cells | N/A |
| | **l** | Representative image | N/A | N/A | N/A |
| | **m** | Heat map | 3 | 152 dividing cells; 258 adjacent nuclei; 273 perpendicular nuclei, 8,621 neighboring nuclei | N/A |

| | | | | | |
|---|---|---|---|---|---|
| | **n** | Distribution comparison | 2 | 200 dividing cells; 369 adjacent cells; 369 perpendicular cells; 12,371 neighboring cells | Tukey's multiple comparison test |
| | **o** | Distribution comparison | 3 | 152 dividing cells; 258 adjacent nuclei; 273 perpendicular nuclei, 8,621 neighboring nuclei | Tukey's multiple comparison test |
| | **p** | Averaged image | 2 | 200 dividing cells; 708 adjacent cells | N/A |
| | **q** | Distribution comparison | 2 | 200 dividing cells; 708 adjacent cells | Paired sample t-test |
| **3** | **a** | Cartoon | N/A | N/A | N/A |
| | **b** | Distribution comparison | 3 | 94 dividing cells | Two-sample t-test |
| | **c** | Distribution comparison | 3 | 300 dividing cells | Two-sample t-test |
| | **d** | Distribution comparison | 3 | 22 dividing cells | Paired sample t-test |
| | **f** | Cartoon | N/A | N/A | N/A |
| | **e, g, h** | Correlation (simulation) | N/A | N/A | N/A |
| | **i** | Cartoon | N/A | N/A | N/A |
| | **j** | Distribution comparison | 2 | 49 control dividing cells; 59 experimental dividing cells | Two-sample t-test |
| | **k** | Distribution comparison | 2 | 20 control dividing cells; 21 experimental dividing cells; 44 control adjacent cells; 47 experimental adjacent cells | Two-sample t-test |
| | **l** | Distribution comparison | 2 | 47 control dividing cells; 59 experimental dividing cells | Two-sample t-test |
| | **m** | Distribution comparison | 2 | 20 dividing cells | Two-sample t-test |
| | **n** | Distribution comparison | 2 | 40 control dividing cells; 40 experimental dividing cells | Two-sample t-test |
| | **o** | Representative image | N/A | N/A | N/A |
| | **p** | Correlation | 2 | 40 dividing cells | Pearson's *r* and linear regression best-fit line |

| | | | | | |
|---|---|---|---|---|---|
| **4** | **a** | Representative image | N/A | N/A | N/A |
| | **b** | Distribution comparison | 3 | 26 dividing cells; 41 adjacent cells; 30 perpendicular cells | Paired sample t-test |
| | **c** | Averaged image | 3 | 26 dividing cells; 41 adjacent cells | N/A |
| | **d** | Distribution comparison | 3 | 26 dividing cells; 41 adjacent cells | Paired sample t-test |
| | **e** | Distribution | N/A | 9 model systems | None |
| | **f** | Distribution | N/A | 9 model systems | None |
| | **g** | Representative image | N/A | N/A | N/A |
| | **h** | Distribution | 2 | 17 dividing cells | One-sample t-test |
| **E1** | | Distribution | 6 | 10 data point measurements averaged per replicate | N/A |
| **E2** | **a, d, g** | Heat map | 3 | 300 dividing cells | N/A |
| | **b, e, h** | Frequency distributions | 3 | 300 dividing cells | N/A |
| | **c, f, i** | Distribution comparison | 3 | 300 dividing cells | Multiple comparison test |
| | **j** | Heat map | 3 | 300 dividing cells | N/A |
| | **k** | Distribution comparison | 3 | 300 dividing cells | Tukey's multiple comparison test |
| **E3** | **a, d, g, j** | Heat map | 2 | 197 dividing cells | N/A |
| | **b, e, h, k** | Frequency distributions | 2 | 197 dividing cells | N/A |
| | **c, f, i, l** | Distribution comparison | 2 | 197 dividing cells | Multiple comparison test |
| **E4** | **a, c, e,** | Heat map | 3 (low, medium density), 2 (high density) | 140 dividing cells (low density), 300 dividing cells (medium density), 168 dividing cells (high density) | N/A |
| | **b, d, f** | Distribution comparison | 3 (low, medium density), 2 (high density) | 140 dividing cells (low density), 300 dividing cells (medium density), 168 dividing cells (high density) | One-way ANOVA post-test for trend |
| | **g** | Representative image | N/A | N/A | N/A |
| | **h** | Correlation | 1 | 3 monolayer samples, 9 fields of view analyzed | Spearman rank correlation |
| **E5** | **a** | Heat map | 1 | 7 dividing cells | N/A |

| | | | | | |
|---|---|---|---|---|---|
| | **b** | Distribution comparison | 1 | 7 dividing cells | Tukey's multiple comparison test |
| **E6** | **a** | Representative image | N/A | N/A | N/A |
| | **b** | Frequency distribution | 3 | 152 dividing cells | $\chi^2$ test against uniform distribution |
| | **c** | Cartoon | N/A | N/A | N/A |
| | **d** | Representative image | N/A | N/A | N/A |
| | **e** | Distribution comparison | 2 | 80 and 67 fields of view analyzed for full-length and truncated E-cadherin samples, respectively. | Two-sample t-test |
| **E7** | **a** | Representative image | N/A | N/A | N/A |
| | **b** | Correlation | 3 | 205 dividing cells | Pearson's *r* |
| | **c** | Distribution comparison | 2 | 46 control dividing cells; 34 experimental dividing cells | Two-sample t-test |
| | **d** | Distribution comparison | 2 | 53 control dividing cells; 54 experimental dividing cells | Two-sample t-test |
| | **e** | Representative image | N/A | N/A | N/A |
| | **f** | Distributions | 2 | 14 dividing cells | One-sample t-test |
| | **g** | Images | 2 | 10 dividing cells | N/A |
| **E8** | **a** | Cartoon | N/A | N/A | N/A |
| | **b-e** | Correlation (simulation) | N/A | N/A | N/A |
| **E9** | **a** | Image | 1 | N/A | N/A |
| | **b** | Representative image | N/A | N/A | N/A |
| **E10** | **a-h** | Single images | N/A | N/A | N/A |

**Supplementary table 2**

List of parameters employed in the computational model.

| Symbol | Definition | Value |
|---|---|---|
| $r_{0,\mathrm{L}}$ | Lower limit of chain length | $5\times10^{-8}$ [m] |
| $r_{0,\mathrm{H}}$ | Upper limit of chain length | $1.25\times10^{-6}$ [m] |
| $\kappa_\mathrm{s}$ | Extensional stiffness of chains | $5.0\times10^{-5}$ [N/m] |
| $\theta_0$ | Equilibrium bending angle formed by adjacent triangles | 0 [rad] |
| $\kappa_\mathrm{b}$ | Bending stiffness of a membrane | $2.77\times10^{-19}$ [N·m] |
| $V_0$ | Equilibrium volume encapsulated by the membrane | $4.19\times10^{-15}$ [$m^3$] |
| $\kappa_\mathrm{v}$ | Strength of volume conservation | 0.07-10,000 [N/$m^2$] |
| $A_0$ | Equilibrium surface area of the membrane | $1.26\times10^{-9}$ [$m^2$] |
| $\kappa_\mathrm{a}$ | Strength of area conservation | $1\times10^{-6}$-$1\times10^{-4}$ [N/m] |
| $\lambda$ | Constant for drag coefficients | $6.44\times10^{-7}$ [kg/$m^2$s] |
| $v_\mathrm{c}$ | Speed of contraction at equator | $3.17\times10^{-8}$ [m/s] |
| $\Delta t$ | Time step | $4.0\times10^{-4}$ [s] |
| $k_\mathrm{B}T$ | Thermal energy | $4.142\times10^{-21}$ [J] |
